# *CDH1* loss remodels gene expression and lineage identity in human mammary epithelial cells

**DOI:** 10.1101/2025.06.20.660633

**Authors:** Maggie Musick, Chinasa A. Ufondu, Carmen E. Rowland, Micaela E. Brown, Joseph L. Sottnik, Madeleine T. Shackleford, Camryn S. Nesiba, Richard Zera, Arthur T. Johnson, Kathryn L. Schwertfeger, Andrew C. Nelson, Julie H. Ostrander, Matthew J. Sikora

## Abstract

Invasive lobular carcinoma (ILC) is a common subtype of breast cancer, molecularly defined by genetic loss of *CDH1,* and subsequent loss of cell adhesion protein E-cadherin, in ∼95% of ILC. Though *CDH1* loss occurs early in ILC oncogenesis, it is unclear how this facilitates transformation. We modeled early *CDH1* loss using “normal” human mammary epithelial cells (HMEC), i.e. finite lifespan cells reflecting early hyperplasia, and targeted E-cadherin signaling using antibodies versus causing genetic *CDH1* loss using siRNA or CRISPR/Cas9-knockout. Transcriptome analysis across four HMEC models showed that the mode of E-cadherin targeting is critical for the subsequent phenotype. Antibody-mediated inhibition of cell-cell contacts induced gene signatures of epithelial-mesenchymal transition (EMT), consistent with the role of E-cadherin suppression during the EMT process. Conversely, genetic *CDH1* loss – as in ILC oncogenesis – repressed EMT signatures, and instead remodeled gene expression toward a luminal epithelial phenotype. RNA-seq, single cell transcriptomics, flow cytometry, microscopy, and ATACseq analyses support that *CDH1* loss induces lineage remodeling to a luminal state, which is mirrored in transcriptomic analysis of clinical ILC precursor lesions. By isolating luminal versus basal cells prior to *CDH1* knockout, we found that *CDH1* loss led to remodeling of lineage identity in both populations, converging on a new lineage homeostasis with a luminal progenitor-like phenotype. Consistent with the shift to a luminal progenitor phenotype, *CDH1* loss enhanced proliferative capacity over the finite lifespan of the HMECs, highlighting a feature of early *CDH1* loss that may contribute to clonal advantage during tumor initiation. Moreover, *CDH1* loss enhanced anoikis resistance, a defining feature of ILC cells. Our findings support that genetic loss of *CDH1* in mammary epithelial cells induces transcriptional and phenotypic changes consistent with lineage identity remodeling toward a luminal progenitor-like state, which may underpin the mechanism by which early *CDH1* loss mediates ILC oncogenesis.

## INTRODUCTION

Invasive lobular carcinoma (ILC) is the most common special histological subtype of breast cancer, accounting for ∼15% of all new diagnoses. ILC is molecularly identified by loss of the cell adhesion protein E-cadherin, encoded by *CDH1*, which in ILC is typically via *CDH1* loss-of-heterozygosity (e.g. *CDH1* loss-of-function mutation with a 16q22.1 deletion) [1]. *CDH1* loss is associated with ILC’s characteristic “single-file” discohesive growth pattern in the breast and unique metastatic spread [2–5], and anoikis resistance, i.e. anchorage-independent survival, another hallmark of ILC [6,7]. Importantly, *CDH1* loss is an early event in ILC oncogenesis found in a majority of ILC precursor lesions; ∼95% of atypical lobular hyperplasia (ALH) and lobular carcinoma *in situ* (LCIS) lesions are E-cadherin-negative [8–11]. However, clinical disease progression in ILC is poorly understood. Though ALH and LCIS are associated with >4-fold and >10-fold increase in breast cancer risk, respectively, clinical management of benign ILC precursors lacks clear guidelines and patients face uncertainty with their long-term risks [9]. Given the central role of *CDH1* in ILC oncogenesis, understanding mechanisms of cancer progression upon *CDH1* loss can improve patient risk assessment and provide new mechanistic insight into the role of *CDH1* as a tumor suppressor.

*CDH1* encodes E-cadherin, a cell surface adhesion protein with critical roles in epithelial cell-cell communication and pathologic contexts in epithelial-mesenchymal transition (EMT), invasion, and metastasis. Though *CDH1* is a tumor suppressor (e.g. hereditary *CDH1* mutations are linked to gastric cancer and ILC [12]), E-cadherin loss or suppression has differing context-specific implications. In breast cancer of no special type (NST; i.e. invasive ductal carcinoma, IDC), E-cadherin is critical for metastasis [13,14]; transient loss of E-cadherin occurs in late stages of cancer progression via epigenetic silencing (e.g. hypermethylation) [15,16], but E-cadherin must be re-activated at metastatic sites to facilitate outgrowth [13,17]. Though E-cadherin suppression is a hallmark of EMT, E-cadherin/*CDH1* loss in ILC is not accompanied by other canonical features of EMT [18], and other studies support that E-cadherin loss leads to induction of EMT specifically in transformed settings [19–21]. Importantly, unlike in EMT, *CDH1* loss in ILC is neither transient nor reversible, thus is mechanistically and phenotypically distinct from EMT-related E-cadherin suppression.

Despite being originally identified in the breast lobules, the cell-of-origin for ILC within the mammary gland has not been explicitly defined. The mammary gland consists of two main epithelial lineages: luminal cells making up the inner lining of the duct, consisting of hormone-responsive and secretory cells, and basal (or myoepithelial) cells lining the duct [22–24]. E-cadherin is expressed in both populations and is critical to maintaining organization in mammary ducts [25,26]. Whether these lineages each differentially support ILC oncogenesis is unclear. However, ILC predominantly present luminal features without classic basal cytokeratins [27,28]) and are largely restricted to luminal-type tumors (i.e. ∼95% of ILC are estrogen receptor α /ER-positive [29]). The potential relevance of lineage is also reflected in genetically engineered mouse models (GEMMs; Cre/l*oxP, Cdh1*^flox/flox^ models) [30], as not all Cre-driving promoters drive ILC-like oncogenesis. Cre driven by the whey acidic protein promoter (*Wap*; mammary secretory/luminal epithelial cells) or the cytokeratin 14 promoter (*Krt14*; broadly active in epithelial cells) both facilitate ILC-like mammary oncogenesis [19,31]. However, Cre driven by the *Lgr6* promoter (active in epithelial stem cells) did not permit ILC-like oncogenesis, instead producing NST/IDC-like mammary tumors with metaplastic squamous features [32]. Taken together, cellular commitment to the epithelial lineage may be important for ILC oncogenesis.

To further examine how genetic loss of *CDH1* facilitates ILC development and functions as a distinct oncogenic event in breast cancer, we modeled different modes of *CDH1*/E-cadherin functional vs genetic suppression in “normal” non-transformed human mammary epithelial cells (HMEC). Strikingly, genetic *CDH1* loss in these models of early hyperplasia drove a distinct form of phenotypic remodeling compared to functional E-cadherin suppression, which we explored further. This approach ultimately points toward a key role for *CDH1* loss in creating a cellular pathway for ILC oncogenesis.

## RESULTS

### Genetic loss of E-cadherin represses EMT and shifts cells to a luminal transcriptional state

To study the role of *CDH1* loss as a driver of ILC oncogenesis, we used finite lifespan human mammary epithelial cells (HMEC) derived from reduction mammoplasties from individual donors. We used variant HMECs that were altered in culture to overcome the senescence barrier (stasis) via overexpression of *CCND1* [33,34]. *CCND1* is a common copy number gain in ILC (∼40% of ILC [35]), thus making these models of early hyperplasia relevant settings for targeting *CDH1* (while facilitating genetic studies, versus using primary cells). Of note, the post-stasis HMEC cultures maintain distinct heterogeneous populations of luminal/basal cells [36]. HMEC models used herein include 122L-D1 (referred to as 122D1), 153L-D1 (153D1), 184D-D1L (184D1), and 240L-D1 (240D1) [36]. We used three modes of E-cadherin suppression across the 4 HMEC models: 1) inhibition of E-cadherin using a monoclonal antibody that binds the extracellular domain of E-cadherin, HECD-1 (α-Ecad); 2) *CDH1* siRNA (si*CDH1*); 3) doxycycline-inducible CRISPR/Cas9 knockout of *CDH1* (*CDH1-*KO) (**Fig. 1A**). For *CDH1*-KO models, we used a non-targeting gRNA (NTG) control and two *CDH1*-targeting gRNA (*CDH1*-KO g1 and g2). *CDH1*-KO was generated by two weeks of Cas9 activation to induce frameshift mutations in *CDH1*, without clonal selection post-knockout to maintain HMEC lineage heterogeneity. In evaluating *CDH1*/E-cadherin targeting, α-Ecad caused HMEC to have a rounded morphology, blocked monolayer formation, and reduced the binding efficiency to E-cadherin-coated plates (Supp. Fig. 1A). siRNA reduced E-cadherin by ∼70% by western blot (Supp. Fig. 1B). CRISPR/Cas9 knockout of *CDH1* was confirmed through flow analysis, complete knockout was observed in up to ∼54% of cells with the remainder showing reduced levels consistent with hemizygous knockout (Supp. Fig. 1C-F).

**Figure 1.**
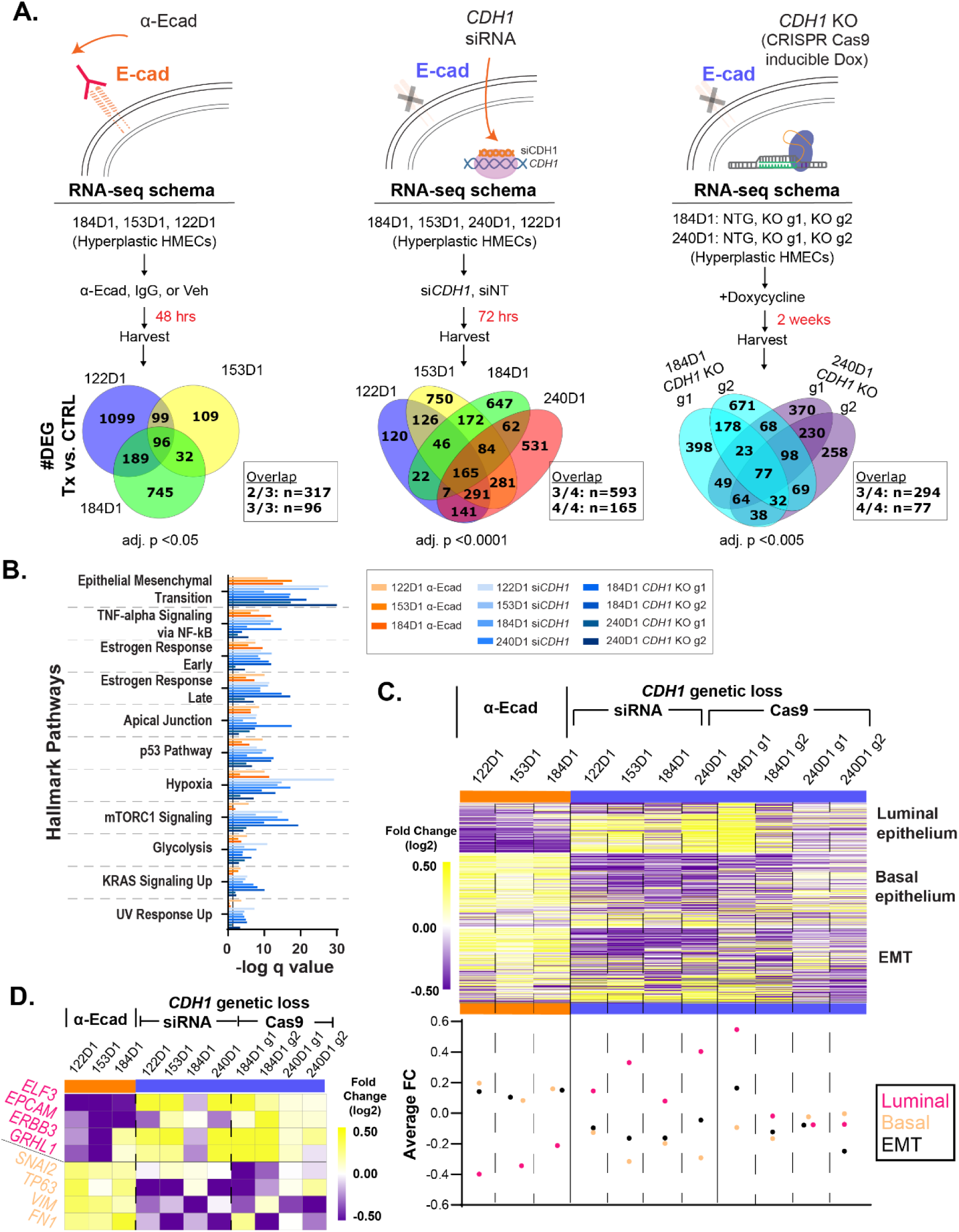
Genetic loss of E-cadherin represses EMT and shifts cells to a luminal transcriptional state. A) Workflow of RNAseq using three methods of E-cadherin suppression; extracellular antibody inhibition (α-Ecad), siRNA transfection (si*CDH1*), and dox-inducible CRISPR/Cas9 (*CDH1*-KO). Venn diagrams show the overlap of differentially expressed genes (DEG), and the adj. p value cutoff used to determine gene lists used for B-E. B) Overrepresentation analysis of Hallmark pathways in all RNA-seq datasets (q value < 0.0001 across genetic loss datasets. Color coded by contact inhibition (orange) vs. genetic loss of *CDH1* (blue). C) Fold change of gene expression (p<0.05) between E-cadherin inhibition and genetic *CDH1* loss within each cell line (HMEC 122L-D1 (122D1), 153L-D1 (153D1), 184D-D1L (184D1), 240L-D1 (240D1)) in the EMT Hallmark, luminal epithelium, and basal epithelium gene signatures. Below heatmap: average fold change of each gene signature across cell lines. D) Fold changes of representative luminal and basal/EMT genes. K-means clusters are represented by the orange and blue bars (Euclidean distance, 2 clusters, max. iterations = 1000).

We performed RNA-seq using all three modes of E-cadherin loss (**Fig. 1A**), which identified 335-1,915 differentially expressed genes per HMEC model and targeting mode (Supp. Table 1). Differentially expressed genes were most strongly enriched in all the models for the Hallmark [37] signature for EMT, and universally enriched for Hallmarks for apical junction signaling, estrogen response, hypoxia, and glycolysis, among others (**Fig. 1B**). We examined these pathways further using fast gene set enrichment analysis (FGSEA) and observed that antibody inhibition of E-cadherin resulted in positive regulation of hypoxia and EMT (Supp. Fig. 2-3), but in contrast, the models of genetic *CDH1* loss showed negative regulation of hypoxia and EMT.

**Figure 2.**
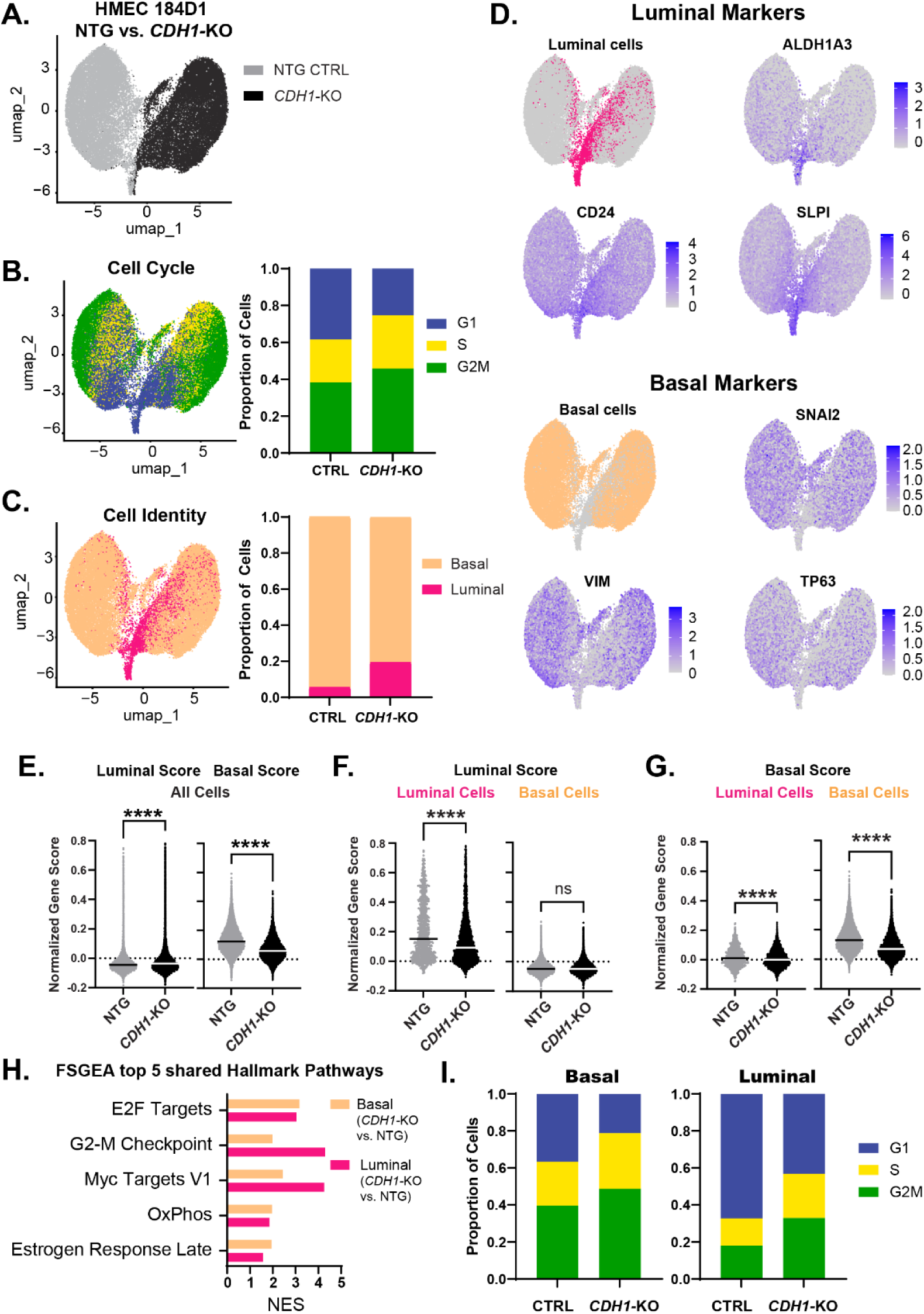
Single cell analysis supports *CDH1*-KO-induced reprogramming in luminal and basal cells. A) HMEC 184D1 *CDH1*-KO g2 (*CDH1*-KO) and HMEC 184D1 NTG UMAP clustering based on sample type. ∼17,000 cells per sample. B) Cells characterized by cell cycle signatures; proportion of cells in each phase by sample type. C) Using z scores of luminal and basal epithelium gene signatures and the AddModuleScore() function in Seurat, cells were classified as luminal (pink) or basal (orange); proportion of cells within each sample type. D) Feature plots of representative luminal (*SLPI, CD24, ALDH1A3*) and basal (*VIM, SNAI2, TP63*) markers. (E-G) The distribution of the normalized gene scores calculated using PIPseeker binning algorithm for both the luminal and basal gene signatures: (E) within all cells (F) luminal scores of the luminal and basal cells (G) basal scores of the luminal and basal cells. H) FGSEA of the top 5 shared Hallmark pathways for luminal (*CDH1*-KO vs. NTG) and basal (*CDH1*-KO vs. NTG). I) Proportion of cells in each cell cycle phase by cell type.-0.3>logFC >0.3, adj. p <1×10^-10^ (pathways p<0.05; except the luminal Estrogen Response Late p=0.055).

**Figure 3.**
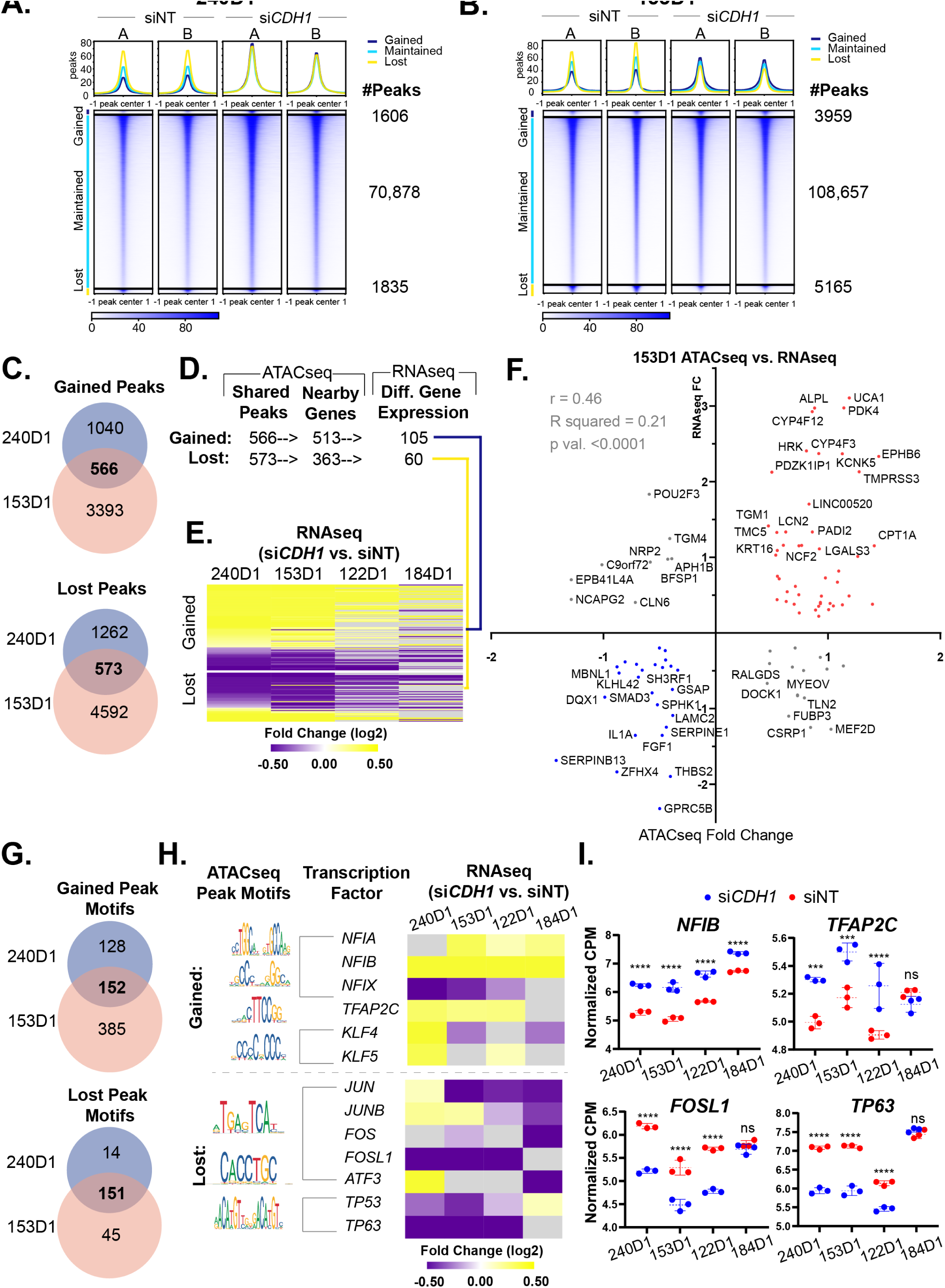
***CDH1***-**KO chromatin landscape supports change in cell identity** A-B) Tornado plots centered at peak centers (+/-1kb) arranged by gained, maintained, and lost peak clusters following si*CDH1*. There are two sets of tornado plots showing the individual replicates of si*CDH1* and siNT for HMEC cell line 240D1 (A) and 153D1 (B). The expression bars below the tornado plots show the relative peak intensity per cell line. Peak plots above each tornado plot show the distribution of peaks per cluster within each sample/replicate. In the HMEC 240D1, there were 74,319 reproducible peaks and of these peaks there were 70,878 maintained peaks, 1,606 gained peaks, and 1,835 lost peaks. In the HMEC 153D1, there were 117,781 reproducible peaks and of these peaks there were 108,657 maintained peaks, 3,959 gained peaks, and 5,165 lost peaks. C) Venn diagrams show the overlap of gained and lost peaks between cell lines. D) Peak to gene analysis (BETA-minus) was used to identify nearby genes within 30kb of the shared peaks. 165 of these genes were differentially expressed in the RNAseq datasets from 240D1 and 153D1 HMEC models using an adj. p < 0.05. E) Heatmaps show log2 fold change (si*CDH1* vs. siNT) of all differentially expressed genes within each of the four HMEC models. Light gray represents non-significant DGE. F) Scatter plot of fold changes from differential peak analysis vs. corresponding gene expression from RNAseq within the 153D1 cell line. Red circles show genes that are upregulated and associated with gained peaks and blue circles show genes that are downregulated and associated with lost peaks. G) ATACseq peak motif analysis (i-cisTarget) was used to identify motifs associated with gained and lost peaks within each cell line (NES > 6 for lost peaks; NES > 3 for gained peaks). Venn diagrams show the overlap of motifs in gained and lost peaks across cell lines. H) Summary of motifs (JASPAR) and corresponding transcription factors organized by motifs associated with gained peaks (top) followed by lost peaks (bottom). Heatmaps show log2 fold change (si*CDH1* vs. siNT) of the transcription factors from the RNAseq datasets of all four HMEC cell lines (adj. p val. < 0.05). I) RNAseq Normalized counts per million (CPM) for each replicate for each condition (siNT, n =3 and siCDH1, n=3) within each HMEC cell model were plotted for the top transcription factors from H. (adj. p val **** < 0.0001; *** < 0.0005).

We further examined changes in EMT and lineage signatures upon *CDH1* loss, using signatures for luminal versus basal epithelial phenotype previously developed through gene expression profiling of these HMEC models (∼300 genes total) [34]. *CDH1* loss by siRNA and CRISPR/Cas9 increased luminal-associated gene expression and decreased basal-associated gene expression, but the opposite was observed with antibody inhibition of E-cadherin (**Fig. 1C-D**). Of note, the apparent modest change in lineage signatures in *CDH1*-KO HMEC 240D1 likely reflects their baseline lineage composition; 240D1 are ∼60% EpCAM^Hi^ luminal cells, whereas 184D1 are >70% EpCAM^Lo^ basal cells (Supp. Fig. 4A). The relative ratio of luminal to basal/EMT genes from Fig. 1D shows that the HMEC 184D1 and 240D1 converge on similar luminal/basal ratios upon Cas9-mediated *CDH1* loss (Supp. Fig. 4B). These data show that genetic loss of *CDH1* in HMEC mirroring the loss of *CDH1* during ILC oncogenesis is distinct from loss of cell-cell contacts by E-cadherin (i.e. by antibody inhibition); the latter activates EMT [21], while the former represses EMT and pushes cells to a luminal transcriptional phenotype.

**Fig. 4.**
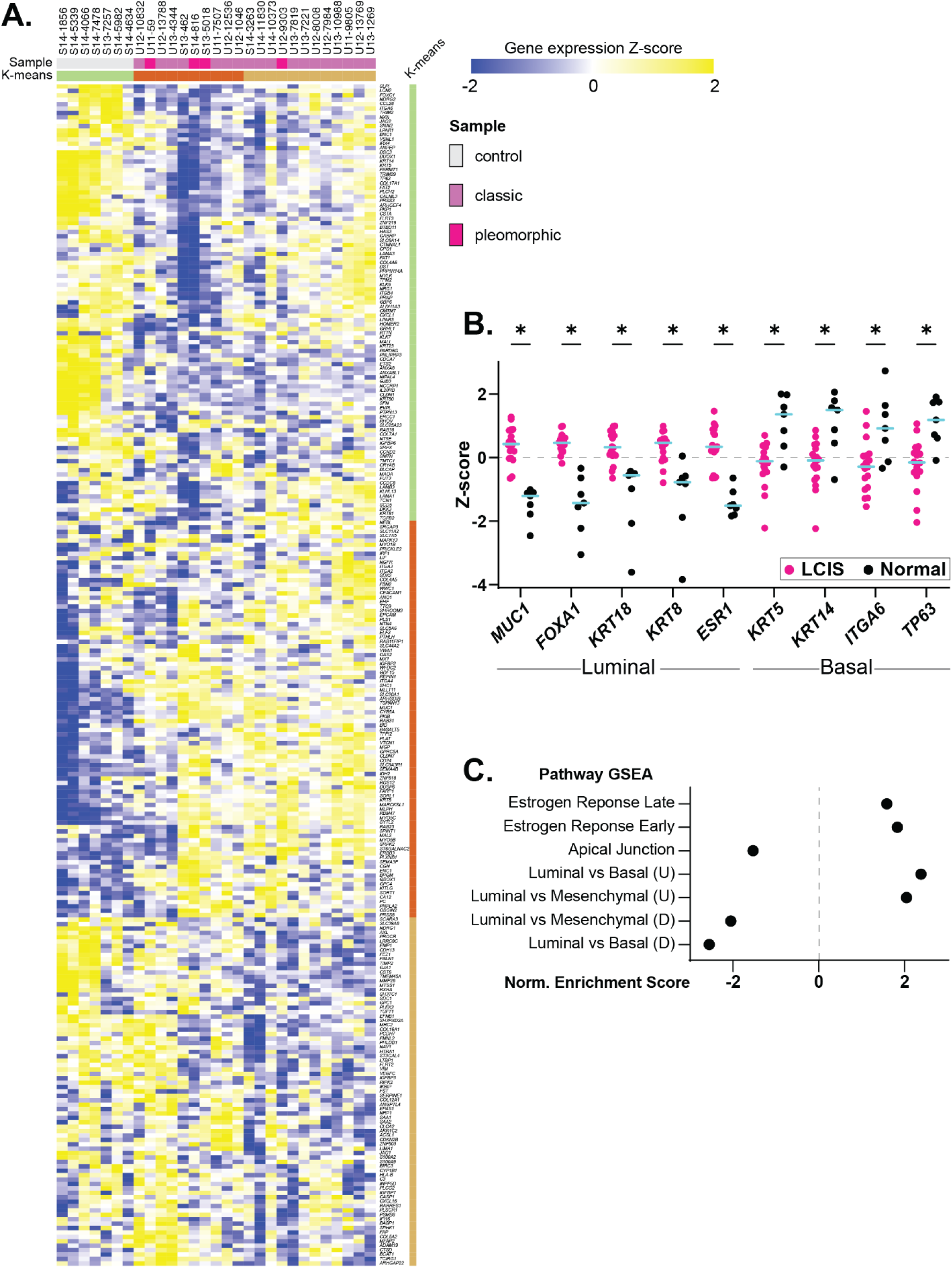
CDH1-KO mimics luminal shift in LCIS data. A) RNA-sequencing on archival tissues from U. Minnesota, from LCIS (n=22) vs. unaffected normal breast from reduction mammoplasties (n=7). Heatmap shows luminal and basal genes and K means clusters (represented by green, orange, and yellow bars). B) Z-scores of key luminal and basal markers comparing LCIS vs. normal breast. *p val. < 0.01. C) Pathway gene set enrichment analysis of Hallmark gene signatures (Estrogen Response Early and Late, Apical Junction) and other gene signatures (Charafe Luminal vs. Basal, Luminal vs. Mesenchymal). Adj. p val. < 0.05.

### Single cell analysis supports CDH1***-****KO*-induced reprogramming in luminal and basal cells

To track the impact of *CDH1* loss within luminal/basal subpopulations in HMEC, we used single-cell RNA sequencing by Particle-templated instant partition sequencing (PIP-seq [38]). We compared HMEC 184D1 NTG versus *CDH1*-KO g2 (referred to as *CDH1*-KO in this analysis), obtaining transcriptome data from 36,956 cells (17,306 NTG, 19,650 *CDH1*-KO), with an average of ∼14,000 transcripts per cell covering ∼3,800 genes per cell (**Fig. 2A**). As in bulk RNA-seq (Fig. 1), the Hallmark EMT signature was repressed across *CDH1*-KO versus control cells (NES=-1.81; adj. p=0.038). Cell cycle state prediction showed an increase in proliferating (S and G2/M) cells in the *CDH1*-KO population (**Fig. 2B**).

We then characterized cells as luminal versus basal using the HMEC signatures described above [34].

*CDH1* loss drove a ∼3.5-fold increase in the luminal population (**Fig. 2C-D**). Accordingly, mean basal signature score decreased across *CDH1*-KO cells while mean luminal signature score increased (**Fig. 2E**). However, changes observed within the luminal and basal subpopulations suggest that *CDH1* loss reduced the fidelity of luminal vs basal lineage identity; i.e. we observed less “polarization” of luminal vs basal features, as indicated by a decreased luminal score in luminal cells (**Fig. 2F**) and a decreased basal score in basal cells (**Fig. 2G**). This reduced lineage fidelity was seen by other mammary cell lineage signatures (Supp. Fig. 5). Other gene expression features were similarly regulated upon *CDH1* loss in both luminal and basal subpopulations (**Fig. 2H**), e.g. Hallmark signatures for oxidative phosphorylation and estrogen response were upregulated in both luminal and basal cells, and there was an increase in proliferative cells (S and G2/M) in both lineage populations (**Fig. 2I**). We isolated cells predicted to be in G1 to exclude these proliferation features; among the most induced genes by *CDH1*-KO were cytokeratin 16 (*KRT16*) and transcription factor inhibitor of DNA binding 2 (*ID2*) (Supp. Fig. 6A-B), both of which are associated with luminal progenitor cells [6,39]. Notably, the cholesterol homeostasis Hallmark was also upregulated in both luminal and basal cells (luminal G1 NES= 2.48, adj. p=0.0027; basal G1 NES= 1.71, adj. p= 0.0706) (Supp. Fig. 6A, C). This single-cell analysis supports observations from bulk RNA-seq that *CDH1* loss drives a population-level shift to a luminal epithelial phenotype, and identifies a loss of strict lineage identity within luminal and basal populations and a gain of features of luminal progenitor cells.

**Figure 5.**
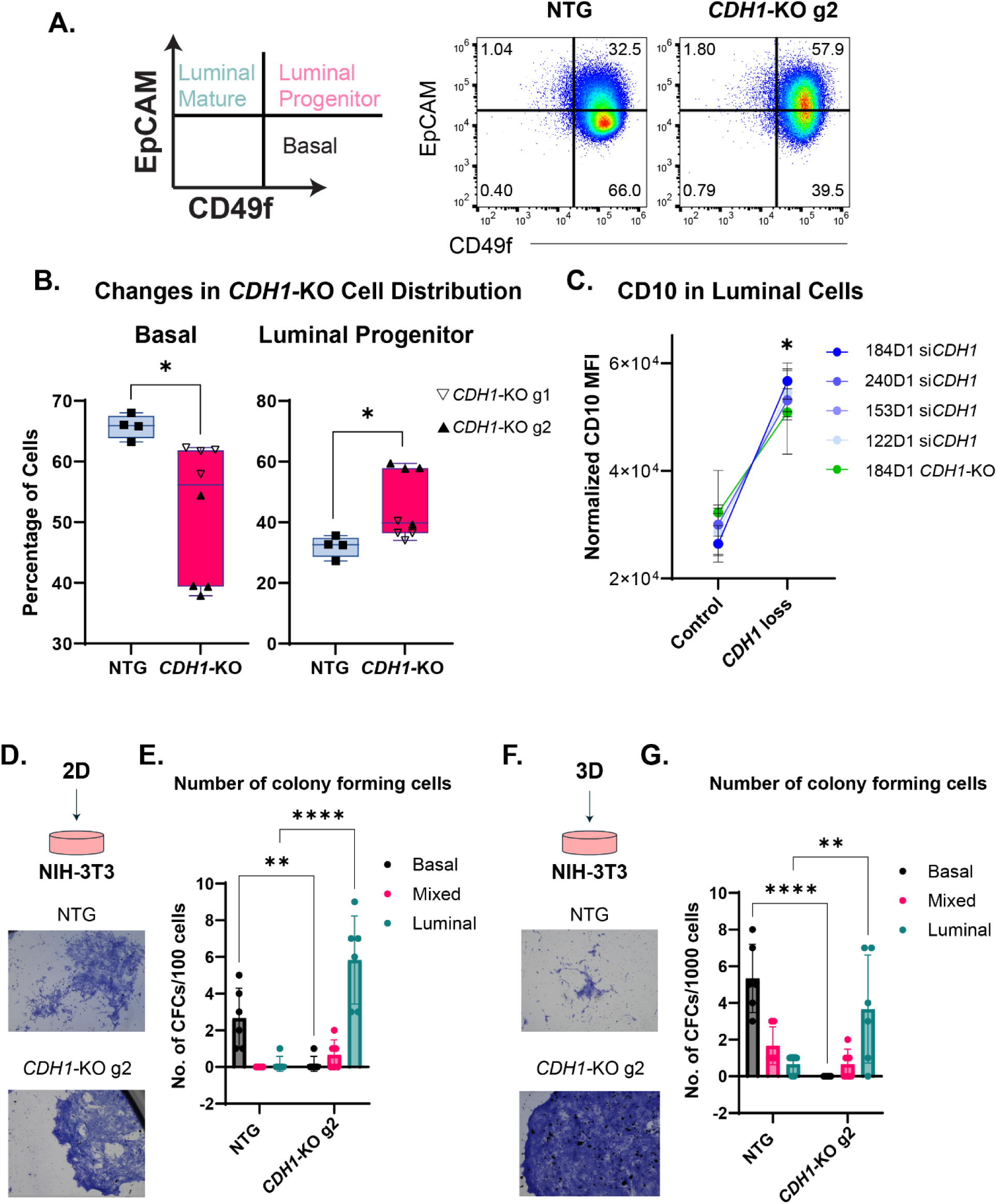
Cellular lineage markers support a *CDH1* loss-induced luminal progenitor-like phenotype. A) Representative plots of flow cytometry analysis of EpCAM/CD49f expression in 184D1 NTG vs. *CDH1*-KO. Gates were set to designate cell lineage: mature luminal (EpCAMhi CD49flow), luminal progenitor (EpCAMhi CD49fhi), and basal (EpCAMlow CD49fhi). Percentages of cells in each lineage are noted in each quadrant. B) Quantification of percentage of basal cells (left) and luminal progenitor cells (right) (n=8; n=4 *CDH1*-KO g1; n=4 *CDH1*-KO g2) *(adj. p val. <0.05). C) Quantification of CD10 expression changes were measured by mean CD10 MFI normalized across multiple experiments (n =14; n= 5 siCDH1 184D1, n=3 si*CDH1* 122D1, n=2 si*CDH1* 240D1, n=4 si*CDH1* 153D1) or *CDH1*-KO vs. NTG (n=8; n=4 *CDH1*-KO g1; n=4 *CDH1*-KO g2) (adj. p val. *<0.0006) D) HMEC *CDH1*-KO g2 and NTG grown in 2D, transferred to feeder layer of irradiated NIH 3T3 cells. Images of NTG and *CDH1*-KO g2 colonies on NIH-3T3 feeder layer E) Quantification of the number of colony forming cells (CFCs) per 100 cells seeded (adj. p val. **** < 0.0001; ** = 0.0017). F) HMEC *CDH1*-KO g2 and NTG grown in 3D, transferred to feeder layer of irradiated NIH 3T3 cells. Images of NTG and *CDH1*-KO g2 colonies on NIH-3T3 feeder layer G) Quantification of the number of colony forming cells (CFCs) per 1000 cells seeded (adj. p val. **** < 0.0001; ** = 0.0020).

**Figure 6.**
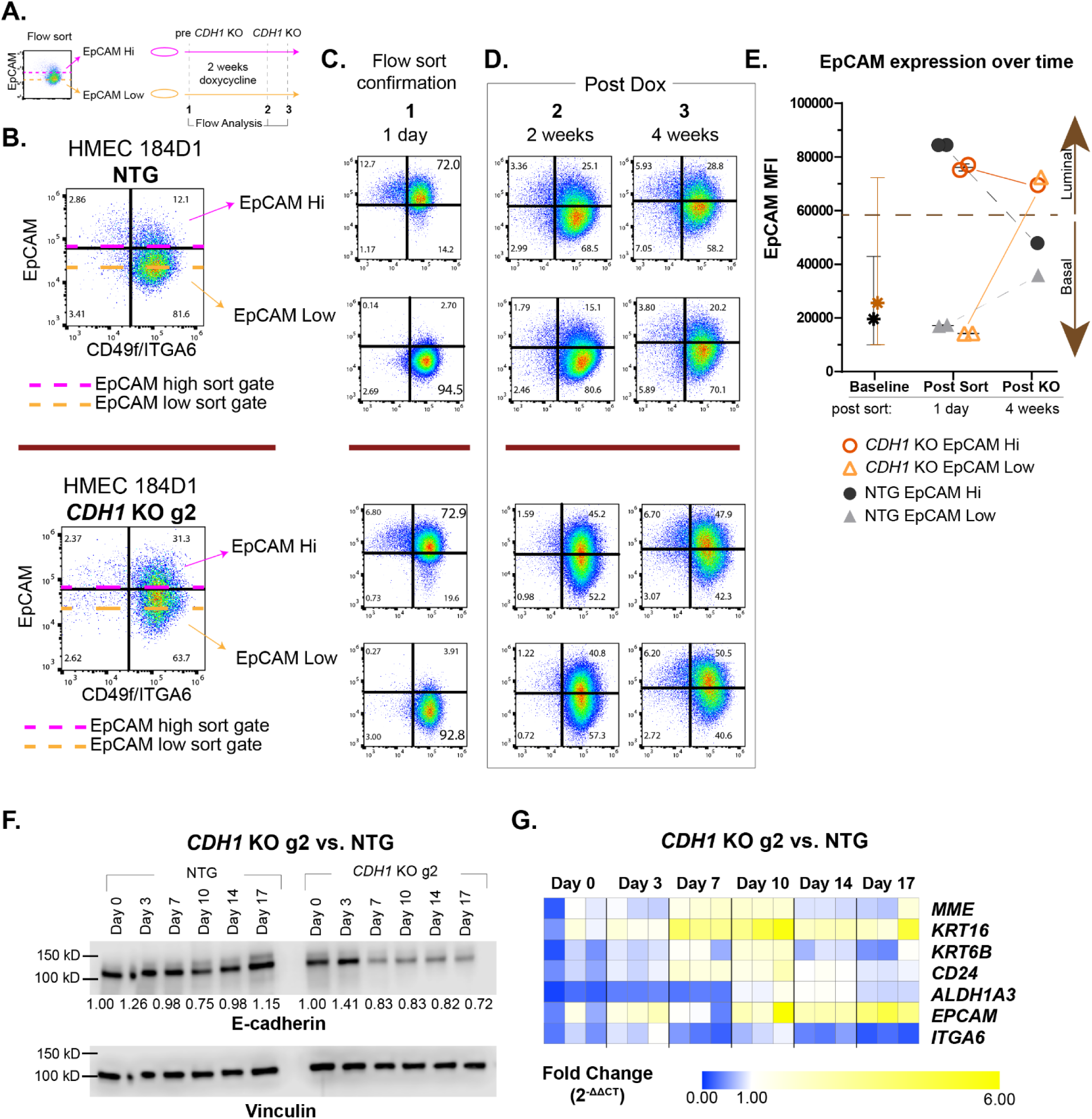
*CDH1*-KO induces lineage remodeling in luminal and basal cells. A) Schematic of flow sorting followed by flow analyses before and after induction of *CDH1*-KO. Cells were sorted into two populations using EpCAM: EpCAM hi (luminal) and EpCAM low (basal) and were separately treated with doxycycline. B) Flow sorting and subsequent flow analyses were conducted in two cell lines: HMEC 184D1 NTG and HMEC 184D1 *CDH1*-KO g2. Dashed lines show where sorting gates were set based off EpCAM expression. C) 1 day post sort (before induction of *CDH1*-KO) the efficiency of the sort was determined by cell distribution based on EpCAM/CD49f expression as described in Fig. 4A (n=1). D) After *CDH1*-KO (2 weeks and 4 weeks post dox induction) flow analysis of cell distribution based on EpCAM/CD49f was determined (n=3; n=2). See gating FMOs for each flow analysis in Supp. Fig. 7. E) EpCAM MFI (geomean) within parental/pre-sorted population (baseline) and each sorted population before induction of *CDH1*-KO (post-sort) and after *CDH1*-KO (post KO). Baseline population error bars show 10% and 90% distribution of EpCAM MFI. F) Time course western blots of E-cadherin with Vinculin loading control. Densitometry was calculated by subtracting local background, normalizing to lane-specific loading control (Vinculin), and then normalized to Day 0 NTG or Day 0 *CDH1* KOg2. G) Paired time course samples analyzed via qPCR for basal and luminal gene expression.

### Chromatin landscape supports CDH1 loss-induced changes in cell identity

Since cell lineage is closely tied to chromatin accessibility [40–42], we examined changes in the chromatin landscape caused by *CDH1* loss via ATAC-seq, using siRNA in HMEC 240D1 and 153D1. The majority of accessible chromatin sites (i.e. peaks) in both models were unchanged by *CDH1* knockdown (**Fig. 3A-B**); gained peaks (chromatin opened by *CDH1* loss) accounted for 2.1% and 6.3% of all peaks in 240D1 and 153D1, respectively, and reduced/lost peaks (chromatin closed by *CDH1* loss) accounted for 2.5% and 8.2% of all peaks in 240D1 and 153D1, respectively (**Fig. 3A-B**). The distribution of maintained and gained peaks were similar in introns (∼40%), distal intergenic regions (∼16%), and promoters (∼37%). However, lost peaks had a higher distribution of peaks found in introns (49-52%) and distal intergenic regions (18-23%) and reduced distribution of peaks found in promoters (18-26%) compared to maintained and gained peak distributions (Supp. Fig. 7A). Between the HMEC models, we see direct overlap of 566 gained peaks and 573 lost peaks (**Fig. 3C**), which were within 30kb of 513 and 363 genes, respectively (**Fig. 3D**), suggesting that these genes were directly regulated by changes in chromatin accessibility upon *CDH1* loss. Accordingly, 165 of the genes identified in ATACseq were differentially expressed after *CDH1* knockdown in both HMEC models (153D1 and 240D1, using a cutoff value of adj. p< 0.05 for differential gene expression) (**Fig. 3D-E**). Genes associated with gained peaks had significantly higher mean fold change compared to the genes associated with lost peaks (Supp. Fig. 7B-C) and included luminal breast cancer-associated markers (*KCNK5, PDK4, CPT1A*)[43–46] (**Fig. 3E-F**, Supp. File 2); genes associated with lost peaks were mostly downregulated and included basal and EMT-related factors such as *SERPINE1*, *SPHK1*, ZFHX4, *LAMC2*, and *NRG1* (**Fig. 3E-F**, Supp. File 2)[47–51]. In both HMEC models, there was a significant positive correlation between fold changes in ATACseq peak signal versus fold changes in corresponding gene expression (**Fig. 3F**, Supp. Fig. 7B). The direct association between chromatin landscape and gene expression supports that *CDH1* loss drives epigenomic reprogramming toward a luminal progenitor-like phenotype.

**Figure 7.**
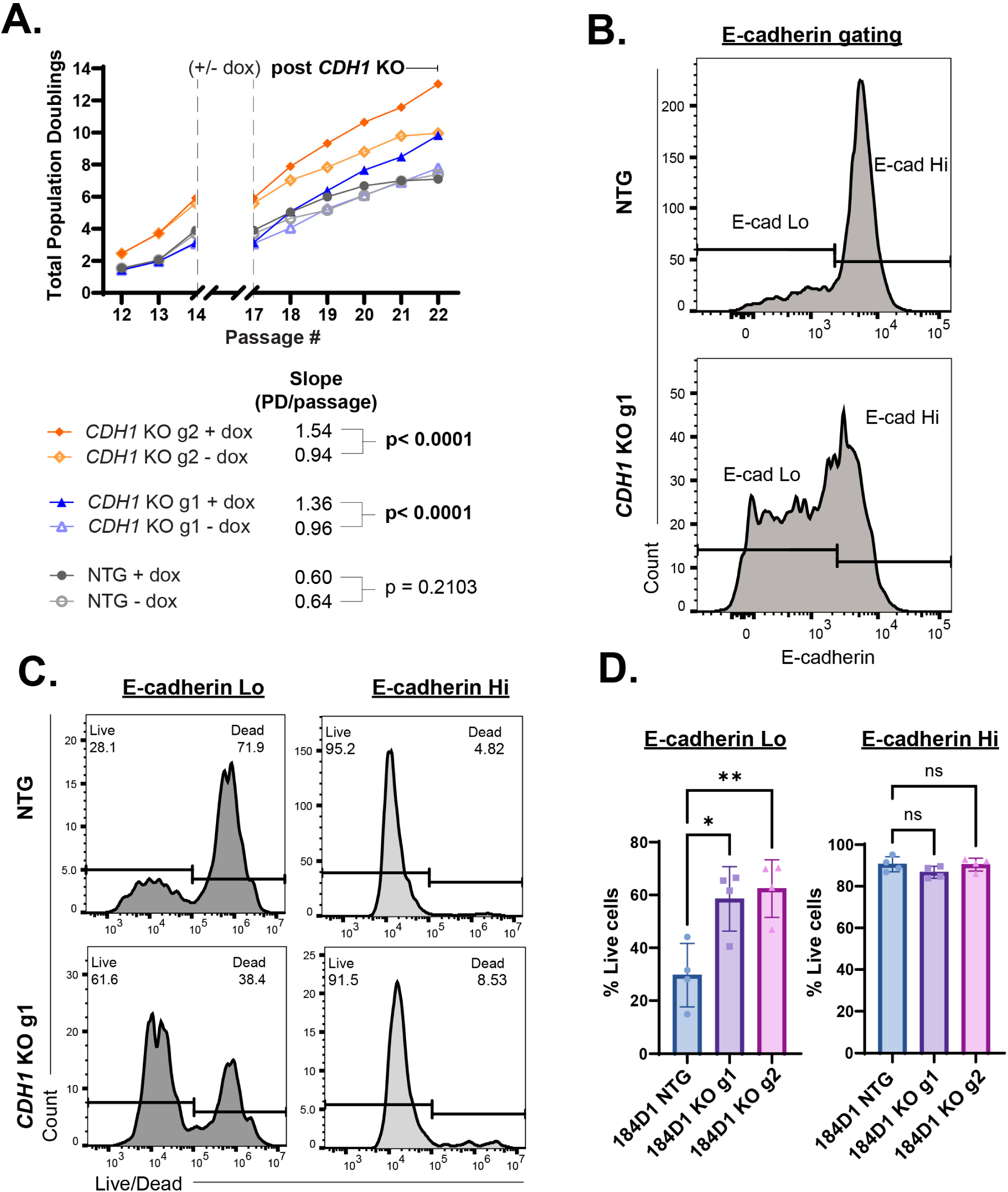
*CDH1*-KO increases proliferative potential and anoikis resistance. A) Total population doublings over the lifespan of the HMEC 184D1 NTG, *CDH1*-KO g1, *CDH1*-KO g2, all +/-dox were calculated by counting cells at the beginning and end of every passage (further described in the methods). The + dox cells were treated from passage 14 through passage 17 with doxycycline (population doublings during this period were excluded from final analysis to remove impact of doxycycline on proliferation). Post-induction proliferation rates were calculated using the following equation: Slope = Δ Population Doublings / 5 passages (17 → 22). Sidak’s multiple comparison test was used to calculate statistical significance between the slopes of the + and – dox models. P value = 0.0126 and 0.0061 for *CDH1*-KO g1 and g2 respectively. P values were calculated using one way ANOVA and Dunnett’s multiple comparisons test. B) Anoikis resistance assay, E-cadherin Lo and Hi gating for NTG and *CDH1*-KO cells. C) Flow cytometry representative histograms of anoikis resistance using live/dead fixable dye in E-cadherin Lo and Ecadherin Hi populations of NTG and *CDH1*-KO cells. D) Quantification of percent live cells as a representation of anoikis resistance in ULA (n=4) in E-cadherin Lo and E-cadherin Hi populations.

To identify potential drivers of the changes in chromatin accessibility after *CDH1* loss, we examined transcription factor motifs enriched in gained versus lost chromatin accessibility sites [52]. There were 152 shared motifs associated with gained peaks and 151 motifs associated with lost peaks (**Fig. 3G**, Supp. File 2). Among the top motif families were NFI-related motifs including (NFIA, NFIB, and NFIX), which are indicative of a luminal progenitor state, ETS-related motif TFAP2C, and epithelial identity motif KLF4 and progenitor-associated motif KLF5 [53–57] (**Fig. 3H**). The gene expression of these factors were mostly upregulated across all four HMEC models following *CDH1* loss; however, there was variability across HMEC models (**Fig. 3H**).

There was no overlap in motifs between gained and lost peaks. The motifs associated with the lost peaks include basal AP1-related motifs JUN, FOS and ATF3 and basal/EMT-related motif TP63 [47,58]. The loss of these motifs in ATACseq were accompanied by downregulation of their transcripts following *CDH1* loss (**Fig. 3H**). Although si*CDH1* in the 184D1 HMECs did not have statistically significant differential gene expression changes for several of the top transcription factors, we found that gene expression (in counts per million) in si*CDH1* 184D1s was similar to si*CDH1* gene expression levels from the other three cell models (**Fig. 3I**).

Further, *TP63* expression decreased in *CDH1*-KO vs. NTG in the 184D1 model (-.20 to-.48 log2 FC). This analysis suggests that shared transcriptional/epigenomic factors are dysregulated across HMECs upon *CDH1* loss.

### CDH1 loss in HMEC mirrors gene expression changes in early lobular neoplasia

We hypothesized that gene expression changes seen in the transition from normal breast tissue to lobular neoplasia, i.e. ILC precursors atypical lobular hyperplasia (ALH) and lobular carcinoma *in situ* (LCIS), would provide a window into the early changes caused by *CDH1* loss, as >95% of ALH/LCIS are already E-cadherin negative [1,8,9,11]. However, LCIS gene expression data are extremely limited, with only a single array-based study reported to date [59]. To examine gene expression changes in LCIS, we conducted transcriptomic analyses of archival LCIS tissues from U. Minnesota (n=22) and unaffected normal breast from reduction mammoplasties (n=7) by RNA-seq (Supplemental File 3). Based on our observations in HMEC, we examined expression of the Luminal/Basal signatures as above and found that these genes drove independent clustering of the normal vs LCIS tissues (**Fig. 4A**). Of note, 4 of 22 LCIS were considered to have pleomorphic histology, and while pleomorphic LCIS clustered with classic LCIS (n=18) by these lineage markers, they were excluded in further analysis. Among lineage markers, key luminal markers (*MUC1, FOXA1, KRT18, KRT8, ESR1*) were increased in LCIS, and basal markers (*KRT5, KRT14, ITGA6, TP63*) were decreased in LCIS compared to normal breast tissue **(Fig. 4B)**. We further examined differentially expressed genes between classic LCIS and normal tissues (n=1455 genes at adj.p < 0.01; Supplemental File 4). By gene set enrichment, the Hallmark signatures for Estrogen Response were the most strongly activated in LCIS, whereas Apical Junction was suppressed, consistent with remodeled cell adhesion and cell surface markers with *CDH1* loss **(Fig. 4C**; Supplemental File 3). Changes in other gene sets for luminal vs. basal / mesenchymal lineage [60] were consistent with a luminal-type cell lineage shift in LCIS relative to normal tissue **(Fig. 4C**; Supplemental File 3). This shift in LCIS contrasts ductal carcinoma *in situ*, which are associated with EMT activation [61]. These data support that early ILC oncogenesis includes a cell lineage shift to a luminal phenotype, as seen in our *CDH1* loss models in HMEC.

### Cellular lineage markers support a CDH1 loss-induced luminal progenitor-like phenotype

We next examined how gene expression changes upon *CDH1* loss would lead to changes in cell surface presentation of lineage marker proteins, thus indicating a lineage phenotype change. Flow cytometry analysis of EpCAM versus CD49f (*ITGA6*) was used to define cells as basal (EpCAM^lo^/CD49f^hi^), luminal progenitor (EpCAM^hi^/CD49f^hi^), or mature luminal (EpCAM^hi^/CD49f^lo^) (**Fig. 5A**). Consistent with the lineage shifts we observed by gene expression, *CDH1* loss caused a shift in 184D1 from primarily basal (EpCAM^lo^) to luminal (EpCAM^hi^) phenotype (**Fig. 5A**), with an enrichment in EpCAM^hi^/CD49f^hi^ cells, indicating an enrichment in the luminal progenitor-like population (**Fig. 5B**). We further examined the luminal enrichment using CD10/*MME*, which is typically a basal marker but also indicates progenitor capacity in EpCAM^hi^ cells [62–64]. Across all HMEC and *CDH1* loss models tested, *CDH1* loss increases CD10 expression in the EpCAM^hi^ luminal population (**Fig. 5C**), supporting that these cells acquire progenitor capacity after *CDH1*-KO. We examined this further using the early common progenitor-derived colonies assay [65], wherein colony formation on a layer of irradiated NIH 3T3 mouse fibroblasts, per colony morphology, can distinguish epithelial progenitor capacity as luminal, basal, or mixed. When inducing Cas9/*CDH1*-KO in standard 2D culture and then transferring to the feeder layer, 94% of NTG colonies presented the basal phenotype, whereas 88% of *CDH1*-KO colonies presented the luminal phenotype (**Fig. 5D-E**). We repeated the experiment by allowing cells to form primary spheres in 3D (ultra-low attachment) after *CDH1*-KO, prior to transfer to the feeder layer, and similarly observed that 70% of NTG colonies were the basal phenotype, versus none in *CDH1*-KO colonies where 85% were the luminal phenotype (**Fig. 5F-G**). Taken together, increased CD10 and the functional shift to luminal phenotype colony formation supports that *CDH1*-KO induces a phenotypic shift to a luminal progenitor-like state.

### CDH1***-****KO* induces lineage remodeling in both luminal and basal cells

Our data suggest that *CDH1* loss drives epigenomic and transcriptomic changes that subsequently drive lineage phenotype remodeling, yet the lineage shift could be explained by a selection against basal cells by *CDH1* loss. To examine this, we use flow sorting to isolate luminal (EpCAM^hi^) vs basal (EpCAM^lo^) cells from 184D1 prior to inducing *CDH1*-KO (**Fig. 6A-B**). Twenty-four hours after initial sorting, we confirmed that cells were effectively sorted into luminal vs basal populations (**Fig. 6C**). We then induced *CDH1*-KO in each sorted population over the course of 2 weeks, reassessing lineage markers at 2 weeks and 4 weeks after initiating *CDH1* knockout (**Fig. 6D**). Flow cytometry gating for each analysis was based on respective fluorescence minus one control (Supp. Fig. 8). 184D1 cells are predominantly basal/EpCAM^lo^ at baseline (∼80%), and both EpCAM^hi^ and EpCAM^lo^ sorted NTG cells re-established this baseline over 2-4 weeks (60-80% basal/ EpCAM^lo^; **Fig. 6B-D, top**). Conversely, in the *CDH1*-KO cells, both EpCAM^hi^ and EpCAM^lo^ sorted cells instead maintained a greater luminal progenitor population (40-60% basal/ EpCAM^lo^; 40-50% EpCAM^hi^/CD49f^hi^; **Fig. 6B-D, bottom**). By 2 weeks post sort, all sorted cells attempted to re-establish HMEC heterogeneity; however, both the EpCAM^hi^ and EpCAM^lo^ *CDH1*-KO populations were more luminal than the NTG EpCAM^hi^ and EpCAM^lo^ sorted populations (Supp. Fig. 9A). By 4 weeks post sort, both sorted populations converged at similar EpCAM levels by mean fluorescent intensity (MFI) for *CDH1*-KO versus NTG models (**Fig. 6E**).

Together these findings support that *CDH1* loss induces lineage remodeling in both luminal and basal cells, towards a new EpCAM^hi^ (luminal) homeostasis, suggesting that *CDH1* loss in either mammary epithelial population may ultimately facilitate ILC oncogenesis.

### CDH1 loss induces a temporal shift in lineage marker expression

Our models of *CDH1* loss include si*CDH1* as an acute *CDH1* loss over 72 hrs, whereas *CDH1*-KO is a more chronic or long-term context, with 2 weeks between inducing *CDH1*-KO and assay, leaving a gap in our phenotypic analyses. We performed a time course analysis of *CDH1*-KO in 184D1, measuring E-cadherin protein and lineage marker gene expression over 17 days upon Cas9 induction. By Day 7 post-induction, E-cadherin protein reached a stable minimum in both *CDH1*-KO gRNA models (**Fig. 6F**, Supp. Fig. 9B).

Decrease in E-cadherin coincided with changes in lineage marker gene expression (**Fig. 6G**). Progenitor markers *MME*/CD10, *KRT16*, and *KRT6B* peaked at 7-10 days **(Fig. 5B**, Supp. Fig. 9C); *KRT16* remained elevated through the time course, consistent with being a specifically luminal progenitor marker [39]. Luminal progenitor marker *ALDH1A3* peaked at 10-14 days and similarly remained elevated versus baseline at 17 days. Luminal marker *CD24* peaked at 7-10 days then decreased, consistent with a transition to a CD24^lo^ progenitor state [66] after 10 days. *EPCAM* (luminal) and *ITGA6*/CD49f (basal) expression mirrored a transition to the luminal phenotype 10-17 days post-Cas9 induction, increasing and decreasing, respectively, to stable points at day 10-14 and beyond (**Fig. 5B**, Supp. Fig. 9C). These temporal changes in epithelial lineage phenotype markers support that early gene expression changes captured with acute *CDH1* loss by siRNA drive a phenotypic shift or remodeling to the luminal progenitor phenotype over the course of 10-17 days.

### CDH1***-****KO* increases proliferative potential and anoikis resistance

We next examined how *CDH1* loss-induced lineage remodeling corresponds to acquisition of cancer phenotypes, focusing on proliferation and anchorage-independent growth (anoikis resistance), to link lineage remodeling to early oncogenesis. First, we examined proliferation over the finite lifespan of our post-stasis HMEC cultures, comparing NTG vs *CDH1*-KO models, +/-2 weeks of doxycycline (dox) to induce Cas9.

*CDH1*-KO cells had more population doublings throughout their lifespan compared to the NTG populations (**Fig. 7A**). Specifically in the post-induction phase (i.e. after doxycycline completion, passages 17–22), *CDH1*-KO g1 and g2 +dox populations showed higher proliferation rates than –dox controls, whereas there was no difference in NTG controls; *CDH1*-KO g1 and g2 models +dox had 2.3-fold and 2.6-fold higher proliferation rates, respectively, versus NTG +dox (**Fig. 7A**). These data support that *CDH1* loss increase proliferative potential in HMEC, which may support oncogenesis, e.g. driving clonal advantage.

One of the defining features of ILC is enhanced anoikis resistance, i.e. anchorage-independent cell survival, which has been mechanistically linked to E-cadherin loss [7]. To assess whether *CDH1* loss was sufficient to support anoikis resistance in HMEC, we induced *CDH1*-KO and then seeded cells into ultra-low attachment plates and assessed cell survival based on E-cadherin status using flow cytometry. A subset of *CDH1*-KO cells remain hemizygous for *CDH1* and E-cadherin positive, while a subset of HMEC as in NTG controls are E-cadherin negative (likely via reduced *CDH1*/E-cadherin in basal cells [67]) **(Fig. 7B)**, so we hypothesized that specifically *CDH1*-KO would permit anoikis resistance in E-cadherin low cells. Among E-cadherin low cells, <30% of NTG cells were viable whereas >60% of *CDH1*-KO cells were viable (i.e. anoikis resistant) **(Fig. 7C-D)**. The enhanced anoikis resistance of E-cadherin-negative *CDH1*-KO cells likely supports early tumorigenesis, and suggests that *CDH1* loss-driven cellular remodeling, and not simply decreased E-cadherin levels, is a critical feature of ILC oncogenesis.

## DISCUSSION

In this study, we sought to better understand the cellular consequences of early *CDH1* loss in ILC oncogenesis. Through transcriptomics, flow cytometry, and colony formation assays, we demonstrated that *CDH1* loss, distinct from E-cadherin inhibition, remodels human mammary epithelial cell lineage identity toward a luminal progenitor-like phenotype. Notably, *CDH1* loss induces lineage remodeling in both luminal and basal cells; either population in 184D1 cells converged on a distinct EpCAM^hi^ state after *CDH1* loss, but NTG controls restored the baseline EpCAM^lo^ state. Gene expression changes were accompanied by *CDH1* loss-driven changes in chromatin accessibility, which suggested putative factors that mediate gene expression changes upon *CDH1* loss. Importantly, gene expression changes we observed in HMEC models were mirrored in clinical samples of LCIS, which presented an increase in luminal markers and decrease in basal markers relative to normal breast tissue, supporting that early *CDH1* loss drives cells to luminal lineage phenotype.

Functionally, *CDH1* loss increased the proliferative capacity and proliferation rate of HMEC cells and facilitated anoikis resistance within E-cadherin-negative cells, linking lineage remodeling to key phenotypes that are likely critical for ILC oncogenesis.

We compared different modes of E-cadherin suppression and hypothesized that loss of cell-cell contacts alone (using E-cadherin antibody, α-Ecad) would not phenocopy loss of *CDH1*, wherein both E-cadherin extracellular contacts and intracellular scaffolding and signaling would be lost. Indeed, these modes of E-cadherin suppression had effectively opposite impacts on gene expression, in particular related to EMT signatures, supporting that *CDH1* loss in ILC is a mechanistically distinct event in breast cancer development and progression. The concept that *CDH1* suppression vs E-cadherin inhibition are mechanistically distinct in mammary epithelial cells was previously observed by the Weinberg Lab [21], reporting that *CDH1* loss (via shRNA) was sufficient to induce EMT whereas a dominant-negative E-cadherin that disrupts adherens junctions was not. Importantly, this prior study was performed in SV40 Large T antigen + TERT immortalized HMLE cells and Ras-transformed derivative HMLER cells, wherein *CDH1* shRNA led to activation of β-catenin which drove EMT. Conversely, in ILC, *CDH1* loss is associated with loss of β-catenin and loss of canonical β-catenin/Wnt signaling [28,68]. Our use of finite lifespan HMEC with only *CCND1* overexpression, rather than immortalized/transformed HMLE and HMLER, may highlight that early loss of *CDH1* prior to the acquisition of extensive oncogenic changes is a critical feature of ILC oncogenesis, e.g. by inducing lineage remodeling that may alter the impact of subsequent genetic changes.

Defining the transcription or epigenomic factors that mediate changes in gene expression and lineage remodeling upon *CDH1* loss is a key next step in understanding the role of *CDH1*/E-cadherin as a tumor suppressor in ILC oncogenesis, and key candidates from our studies include inhibitor of DNA binding 2 (ID2). ID2 regulates commitment to luminal lineage [69], and has been shown to be transcriptionally induced upon E-cadherin loss whereupon it drives anoikis resistance in multiple models of ILC [6]. However, anoikis resistance and cell cycle regulation by *CDH1* loss-induced cytoplasmic ID2 is well studied, but the role of nuclear ID2 is less well understood [6]. ID2 does not directly bind to DNA, instead binding other transcription factors including basic helix-loop-helix (bHLH) and ETS family members [70]. In our ATACseq analyses, gained peaks upon *CDH1* loss in both 153D1 and 240D1 are strongly associated with bHLH and ETS motifs including TFAP2C, ELK4, and GABPA. Notably, lost/decreased motifs also have a connection with ID2, including TCF12 and p63. TCF12 can be inhibited by direct ID2 binding in lymphoid progenitors [71], and p63 has been shown to repress *ID2* transcription via direct promoter binding [72]. Of note, *CDH1* loss-driven reduced accessibility at p63 motifs is consistent with the role of p63 as a basal transcription factor [73]. Taken together, changes in chromatin accessibility and associated gene expression point toward ID2 as a potential driving factor of *CDH1* loss luminal reprogramming, with the connection between *CDH1* and ID2 also representing an important area of investigation.

We defined the putative lineage progenitor phenotype in *CDH1*-loss HMEC through upregulation and cell surface presentation of key markers (e.g. EpCAM^hi^ / CD49f^hi^ / CD10^hi^ status), and functional assays including the progenitor-derived colony forming assay, though historically, progenitor populations have been inconsistently defined. However, recent advances, such as single cell RNAseq of human mammary cells compiled into the human breast cell atlas, have identified a broad diversity of cell types within the luminal and basal compartments within and across individuals [74]. Several luminal progenitor populations based on unbiased clustering have been defined by cytokeratin composition and expression of specific transcription factors. One such cluster (LASP2) was defined by increased expression of *KRT16* and *KRT6B*, which we observed in our *CDH1* loss models. The LASP2 cluster was also associated with reduced lineage fidelity, mirroring our observations. This KRT16+ *CDH1* loss-induced progenitor population could be distinctly susceptible to supporting progression toward ILC. Defining the subsequent susceptibility, and how *CDH1* loss interacts with subsequent genetic hits in the path to transformation, is important to understanding ILC oncogenesis. The HMEC models we used represent one stage of a series that have been moved through stepwise genetic progression from primary cells to transformed tumorigenic cells [36], and as such, make a strong platform to apply additional genetic hits to continue beyond early hyperplasia with *CDH1* loss, to overt ILC.

*CDH1* loss induces luminal progenitor-like reprogramming in luminal and basal mammary cells, creating a cellular context that may be permissive to ILC oncogenesis. By identifying potential mechanistic drivers and proliferative processes associated with this reprogramming, our findings highlight new opportunities for identifying, profiling, and targeting early lobular lesions to assess risk and prevent progression to ILC. This work establishes a foundation for future investigations into the etiology of ILC, including mechanisms underlying its distinct patterns of invasion and metastasis.

## MATERIALS AND METHODS

### Cell line culture and conditions

Female post-stasis human mammary epithelial cells (HMEC 184D-D1L 122L-D1, 153L-D1, 240LB-D1) (provided by Dr. Martha Stampfer, Lawrence Berkeley National Lab; https://hmec.lbl.gov) were maintained in 1:1 MEBM:DMEM/F12. MEBM (Lonza, cat#cc-3151) + 5.0ug/mL Insulin (Thermo Fisher, cat# 41400-045), 70.0ug/mL bovine pituitary extract (Thermo Fisher, CAT # 13028014), 0.5ug/mL hydrocortisone (Sigma-Aldrich, Cat# H4001-1G), 5.0ng/mL EGF (Fisher Scientific, Cat # 236EG200), 5.0ug/mL transferrin (Invitrogen, cat# 11108-016), 10^-5^M Isoproterenol (IP) (Sigma, CAT# I5627), and 2.0mM glutamine (Sigma, Cat# 59202C). DMEM/F12 (Corning, CAT # 10-092-CV) + 10ug/mL Insulin, 0.1ug/mL hydrocortisone, 2.5mL BSA (Sigma, CAT# A4161), 0.005ug/mL EGF, 2mM glutamine, 10ug/mL Cholera Toxin (Sigma, CAT #C8052), 1uM Oxytocin (Sigma, CAT# O6379-1MG), and 5% Albumax (Invitrogen, CAT # 11020-021). Cells were incubated at 37°C in 5% CO2. All lines were confirmed Mycoplasma negative and authenticated by STR profiling annually; validated cultures were maintained for no more than 3 months before thawing new cultures.

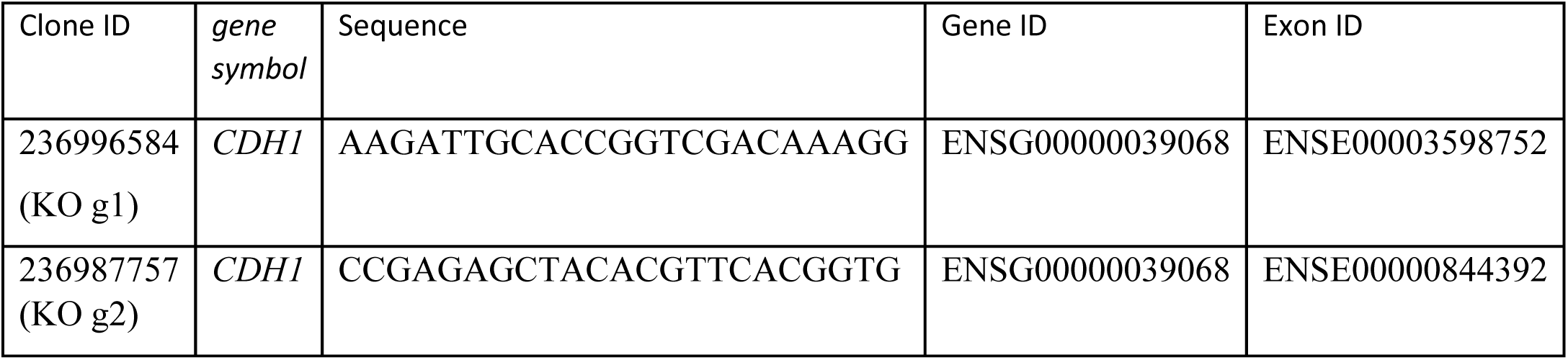

### E-cadherin antibody inhibition

E-cadherin inhibition was performed using HECD-1 antibody (Thermo Fisher, CAT# 131700) according to manufacturer’s instructions. For RNA-seq, 184D-D1L 122L-D1, 153L-D1 HMEC were plated in 48 well plates and left overnight. Cells were treated with HECD-1 or IgG (whole mouse) at 25ug/mL for 48 hours. Cells were washed twice with PBS and stored in-80°C until RNA extraction.

### siRNA knockdown protocol

siRNA was reverse transfected using RNAiMAX (Thermo Fisher) by manufacturer instructions. Constructs are siGENOME SMART pool siRNAs (GE Healthcare Dharmacon): nontargeting pool #2 (D-001206-14-05) and human *CDH1* pool (M-003877-02-0005).

### Inducible CRISPR/Cas9

CRISPR/Cas9 models were established using Cas9 lentiviral vectors and lentiviral vectors obtained from CU Anschutz Functional Genomics Facility (1 control +2 *CDH1*-targeting gRNA; lentiCRISPRv2-puro). Mutation found in KO g2 is truncating at exon 2 and KO g1 is truncating at exon 3. These exons make up the propeptide/signal domain, where one of the most commonly recurring *CDH1* mutations in ILC can be found [75].

Induction of Cas9 and gRNAs was conducted over a 2-week period with fresh doxycycline (1μg/mL) every three-four days. To preserve HMEC line heterogeneity, a mixed population workflow was followed (i.e. cells were not clonally selected after doxycycline induction).

### RNA isolation

RNA extractions were performed using the RNeasy mini kit (Qiagen, cat# 74034); mRNA was converted to cDNA on an Eppendorf Mastercycler Pro (Eppendorf, Hamburg, Germany) and using iSCRIPT cDNA synthesis kit (BioRad, CAT# 1708890).

### RNA-seq

Library preparation and sequencing was performed at the U. Colorado Comprehensive Cancer Center Genomics Core Facility. Libraries were generated with Illumina Stranded mRNAPrep, and sequenced on a NovaSEQ6000 with 2150 paired-end reads, targeting 40106 clusters/80106 reads/sample (final mean= 39,019,637 clusters/sample). Data were analyzed with the UCCC Biostatistics and Bioinformatics Shared Resource.

Illumina adapters and the first 12 base pairs/read were trimmed using BBDuk (RRID:SCR_016969) and<50 bp reads post-trimming were discarded. Reads were aligned and quantified using STAR (2.6.0a, RRID:SCR_004463) against the Ensembl human transcriptome (hg38.p12 genome,release 96), then normalized to counts per million by edgeR package (RRID:SCR_012802). Differential expression was calculated using limma R package (RRID:SCR_010943) and voom() function. Overrepresentation analysis was performed with cluster Profiler R (RRID:SCR_016884) and Molecular Signatures Database (RRID:SCR_016863) gene sets.

### RNA-seq: quantification and statistical analysis

RNAseq was performed using biological replicates, n=3. All cells for each cell line were plated in the same 48 well plate. Biological triplicates are defined as 3 separate wells within the same plate. Reads were aligned and quantified using STAR (2.6.0a, RRID:SCR_004463) against the Ensembl human transcriptome (hg38.p12 genome,release 96), then normalized to counts per million by edgeR package (RRID:SCR_012802).

Differential expression was calculated using limma R package (RRID:SCR_010943) and voom() function. Overrepresentation analysis was performed with cluster Profiler R (RRID:SCR_016884) and Molecular Signatures Database (RRID:SCR_016863) gene sets. For antibody inhibition: differential gene expression was analyzed with gene list cutoff using the adjusted p value 0.05. DGE lists were identified following comparison of DGE between HECD-1 vs. CTRL and IgG vs. CTRL, the final list included DGE only from HECD-1. This indirect comparison was performed, rather than direct HECD-1 vs. IgG (negative control) due to a higher than expected immune response induced by the IgG. For all RNA-seq datasets: enriched pathways were identified using Enrichr. Heatmaps were generated using Morpheus with marker selection-signal to noise (α-Ecad vs. si*CDH1* and *CDH1*-KO).

### Single cell RNA-seq

Following CRISPR editing, single cell suspensions were prepared according to the recommendations of the Illumina Single Cell 3’ RNA Prep, T20 kit (20135692). A detailed workflow of cell processing, mRNA capture, and library preparation can be accessed at https://www.fluentbio.com/resource-category/user-guides/. Samples were pooled and sequenced on a 375Gb lane of the NovaSeq XPlus 10B platform using 150 paired-end base pair reads. Samples were demultiplexed and transcript mapping was performed using PIPseeker software, available at https://www.fluentbio.com/products/pipseeker-software-for-data-analysis/, yielding 1.1 billion barcoded reads between the two samples. Data was processed and analyzed using the Seurat R package [76].

High quality cells were filtered based on transcript count, gene count, mitochondrial content, and genes/UMI metrics. Prior to downstream analysis gene expression data was normalized and scaled. The top 3,000 variable genes were identified and a PCA and UMAP dimensionality reduction were performed. Cell clusters were defined using nearest-neighbor graph analysis and Seurat clustering algorithms. Cells were annotated as luminal or basal using established lineage marker gene signatures and module scoring. Independent reclustering of luminal and basal cell populations were performed. More detailed methods can be found in supplemental methods.

### ATAC-seq

For ATACseq, cells (HMEC 240D1 and 153D1) were plated in 6 well plates and treated with siRNA (siNT or si*CDH1*) for 72 hours. Cells were harvested, counted, and stored at-80C following Active Motif ATACseq cell preparation protocol. Cells were thawed, counted, *Drosophila* Spike-In control (Active Motif #53154) was added, and we proceeded to follow Active Motif’s ATAC-seq protocol (#53150). Transposed DNA was indexed using Illumina’s Nextera adapters within Active Motif’s ATAC-seq kit (#53150). Samples were checked for quality and sequenced by Novogene. Samples were sequenced with 20M PE reads on 10B NovaSeqXPlus.

### ATAC-seq data analysis

Sequence quality was assessed using FastQC (v0.11.9) and MultiQC (v1.14). Reads were trimmed for Nextera adapters and low-quality bases using BBDuk (v39.01), followed by contamination screening with FastQ Screen (v0.15.3). Alignments to the GRCh38 human reference genome were performed using Bowtie2 (v2.5.0). Post-alignment, mitochondrial reads were removed, and fragments were filtered to retain high-confidence, properly paired alignments (MAPQ > 30) using Samtools (v1.16.1). Artifact-prone regions were excluded using the GRCh38 unified blacklist and Bedtools (v2.29.1). For comparative analysis, libraries were normalized by randomly sub-sampling reads to a uniform depth across samples. Normalized coverage tracks (BigWig) were generated with deepTools bamCoverage (v3.5.2). Peaks were called using MACS3 (v3.0.3), identifying high-confidence narrow peaks with a significance threshold of q < 0.05. Data were visualized using the Integrative Genomics Viewer (IGV, v2.18.4).

Differential accessibility analysis was performed using DiffBind (v3.20), retaining only reproducible peaks present in at least two samples. Sub-sampled reads within these consensus regions were quantified via summarizeOverlaps, normalized using Relative Log Expression (RLE) scaling based on full library sizes, and modeled with DESeq2 to identify significant differentially accessible regions (FDR < 0.05, minimum 1.25-fold change) between siCDH1 and siNT conditions. Genomic distribution of peaks (maintained, gained and lost) was determined using R/Bioconductor package ChIPseeker [77].

BETA-minus was used to identify nearby genes within 30kB of a peak [78]. BETA-basic was used to perform an integrated analysis of ATACseq and RNAseq from HMEC 153D1 and HMEC 240D1 cell models (Web-based platform Cistrome). Heatmaps were created using “compute matrix” and “plotHeatmap” tools from web-based platform Galaxy (see supplemental methods for more details).

Motif enrichment analysis was performed using i-cisTarget. First, we did individual analyses using the BED files of gained peaks and lost peaks for both cell lines. Then we used i-cisTarget comparative analysis to find the shared motifs within the gained peaks across both cell lines and within the lost peaks across both cell lines.

### LCIS sample selection, pathology review, RNA isolation, and bioinformatics

*Sample Selection and Pathology Review:* IRB approvals were obtained from both participating institutions (University of Minnesota Study # 1406E51485, July 05, 2014; Hennepin County Medical Center Study # 14-3842, July 9, 2014). A total of 198 cases that contained “LCIS” in the diagnostic report were identified between the years 2010 – 2014. Pathology slide review identified 24 cases of cLCIS and 7 cases of potential pLCIS which contained sufficient LCIS tissue for analysis (>20 mm2). LCIS lesions were adequately separate from any concomitant invasive cancer to ensure accurate enrichment of only LCIS from FFPE sections. Cases were independently reviewed by 3 board-certified surgical pathologists (MK, SC, ACN) for diagnostic classification as classic or pleomorphic LCIS; only cases with independent, 3-sided consensus agreement were classified as pLCIS and all others were classified as classic. One pathologist evaluated the cohort for 6 additional qualitative morphologic characteristics: nuclear contour, nuclear size, chromatin quality, extent of necrosis, presence of mitotic figures, and inflammatory infiltrate (scored on a 1-3 scale). For necrosis scoring: absent = 1; rare, small foci < 5 cell diameters = 2; and prominent central necrosis = 3. For mitoses scoring: absent to rare mitotic figures across all involved lobules = 1; the presence of at least 1 mitotic figure in two or more involved lobular units = 2; and the presence of multiple lobular units with >1 mitotic figure each = 3. Neoplastic cell percentage was minimally 50% in areas macro-enriched for analysis; cases were further categorized into three tiers of neoplastic cellularity: 50-60%, 60-80%, and >80%. 7 control cases from mammoplasty reduction were identified to obtain benign breast tissue.

*FFPE RNA isolation:* Four consecutive 7-µm thick section of FFPE breast tissue from each FFPE block were cut using a fresh microtome blade and mounted on new RNA-free slides. Equipment sterilization with RNase AWAY (Molecular BioProducts Inc, San Diego, CA, USA) was completed between blocks. Following mounting, sections were immediately transferred to a-80C freezer and stored no longer than 2 weeks. Xylene deparaffinization and staining were completed as described in the Arcturus Paradise PLUS reagent system user guide (Life Technologies, Carlsbad, CA, USA). Areas containing cLCIS, pLCIS and representative normal breast tissue were identified and macro-dissected and stored at-80C. Total RNA was then isolated and purified using the Qiagen’s RNeasy Mini Kit (Hilden, Germany). RNA concentration and purity were measured with a Nano-Drop 2000 spectrophotometer (Thermo FisherScientific, Waltham, MA, USA).

RNA Quality Control and NGS Sequencing: RNA-seq was preformed using the TruSeq RNA Access Library Prep Kit from Illumina (San Diego, CA, USA). RNA sample quantity and quality was assessed using a 2100 Bioanalyzer (Agilent, Santa Clara, CA, USA) following the DV200 recommendations of the TruSeq RNA Access Kit. Sequencing was completed on an Illumina HiSeq 2500 with 2 x 100 bp reads, producing an average of approximately 27 M reads/sample. Successful RNAseq data were obtained from 19 cLCIS, 3 pLCIS, and 7 control samples. FPKM counts for all samples are included as Supplemental Data File 1.

*Bioinformatic and Statistical Analysis:* Reads were aligned and quantified using STAR (2.6.0a, RRID:SCR_004463) against the Ensembl human transcriptome (hg38.p12 genome,release 96), then normalized to counts per million by edgeR package (RRID:SCR_012802). Differential expression was calculated using limma R package (RRID:SCR_010943) and voom() function. Heatmap was created using Morpheus (https://software.broadinstitute.org/morpheus) and clusters were defined using K means. Gene set enrichment analysis and overrepresentation analyses were performed using BBSR tools at CU Anschutz (https://bioinformatics.cuanschutz.edu/).

### Flow Cytometry

For extracellular flow cytometry, cells were harvested with 0.05% trypsin/EDTA. Cells were counted, washed with PBS, and resuspended in staining solution (PBS, 1% BSA, 1% EDTA). Cells were plated into Falcon 96-well round bottom plates (Corning # 353077) at 400,000 cells/well. EpCAM-BV421 (Biolegend #324220, clone 9C4, mouse IgG2b, 1:200), CD10-APC/Fire 750 (Biolegend #312230, clone HI10a, mouse IgG1, 1:200), and CD49f-PE (Biolegend #313612, clone GoH3, Rat IgG2a, 1:400) were added to cells in staining solution for 15 minutes on ice. Cells were centrifuged for 3 min at 1400rpm. Staining solution was aspirated and cells were washed with PBS for 10 min on ice. Cells were centrifuged for 4 min at 1400rpm. PBS was aspirated and cells were resuspended in flow solution (PBS, 2% BSA, 0.2% EDTA). Cells were then analyzed in a Novocyte Penteon Flow Cytometer. SpectraComp beads (Bio Slingshot # SSB-05-B), unstained cells, and FMOs were used as controls.

For flow sorting, cells were prepared following the same protocol as the extracellular flow cytometry analysis. Cells were only stained with EpCAM-BV421 (Biolegend #324220, clone 9C4, mouse IgG2b, 1:200). Cells were sorted using the Sony MA900. Gates were set using unstained controls.

For intracellular flow cytometry, cells were harvested with 0.05% trypsin/EDTA. Cells were counted, washed with PBS, and fixed in 4%PFA (100uL/1×10^6^ cells) and incubated at room temperature for 8 minutes. Cells were washed with PBS by centrifugation. Cells were resuspended in 1% Triton-X (100uL/1×10^6^ cells) and incubated at room temperature for 10 minutes. Cells were then washed with PBS by centrifugation and resuspended in staining solution. E-cadherin-AF647 (Cell Signaling #9835, clone 24E10, Rabbit IgG, 1:200) was added to the cells and the staining flow protocol above was followed.

### Population Doubling Assay

Total population doubling over a finite lifespan for each culture was calculated using the formula: population doubling= log_2_(N_final_/N_initial_), where N_initial_ is the number of cells seeded in a dish at each passage and N_final_ is the number of cells recovered from the dish (adapted from Garbe et al. 2009).

### Primary Sphere Culture and Fibroblast Co-culture Assay

Primary mammary epithelial cells were cultured in 2D or as spheres using ultra-low attachment conditions. Briefly, 1 × 10⁵ cells per well were plated in 6-well ultra-low attachment plates and maintained in M87A medium for 7 days to allow sphere formation. Spheres were collected into 2-mL microcentrifuge tubes and pelleted by centrifugation at 900 rpm for 5 minutes at room temperature. The supernatant was aspirated, and sphere pellets were resuspended in 0.5–1 mL of 0.05% trypsin/0.025% EDTA, depending on pellet size, and incubated at 37°C for 10 minutes for enzymatic dissociation. Mechanical dissociation was performed by gentle pipetting (5–10 times) using a 1000-μL pipette. The resulting cell suspension was transferred to 15-mL conical tubes, and trypsin was neutralized with 5–10 mL of calcium-and magnesium-free PBS (CMF-PBS) supplemented with 10% fetal bovine serum (FBS). Cells were passed through a 40-μm nylon mesh filter, pelleted by centrifugation (900 rpm, 5 minutes, room temperature), and resuspended for viable cell counting using a hemocytometer.

For co-culture experiments, NIH-3T3 murine fibroblasts were irradiated with 30 Gy one day prior to use. Irradiated fibroblasts were plated in 24-well tissue culture plates at a density of 2 × 10⁴ cells/cm² in DMEM supplemented with 10% FBS. On the day of sphere dissociation, single-cell suspensions derived from primary spheres were seeded onto the NIH-3T3 feeder layer at a density of 100 cells/mL for 2D culture and 1,000 cells/mL for 3D culture. Co-cultures were maintained at 37°C in a humidified incubator.

Medium was replenished every 48 hours by performing a half-medium change, in which approximately half of the conditioned medium was removed and replaced with fresh M87A medium. After 10 days of co-culture, colonies were gently washed with 0.5–1 mL CMF-PBS and fixed with 250–500 μL ice-cold methanol for 5 minutes, taking care to avoid direct disruption of colonies. Methanol was aspirated, and colonies were stained with Wright dye (150–200 μL per well) for 1 minute at room temperature prior to downstream analysis.

### Anoikis resistance assay

*CDH1*-KO was induced in HMEC 184D1 (NTG used as control). Cells were harvested, resuspended, and transferred to ultra-low attachment plates for 3 days. Cells were harvested, washed, and resuspended in 1xPBS with fixable live/dead dye according to the protocol (Thermo Fisher # L23105). Then cells were fixed and prepared for flow cytometry as described above. Cells were also stained with E-cadherin-AF647 (Cell Signaling #9835, clone 24E10, Rabbit IgG, 1:200). Unstained samples (mix of NTG and *CDH1*-KO cells) were used to set the gate for E-cadherin Hi and E-cadherin Lo populations. Anoikis was quantified based on live/dead fluorescence in E-cadherin Hi and E-cadherin Lo populations, separately.

## ACKNOWLEDGEMENTS

We thank Martha Stampfer, Jim Garbe, and Mark LaBarge for the HMEC cell lines and their invaluable guidance and feedback throughout this project.

This study was partly supported by the National Institutes of Health P30CA046934 by utilizing the Bioinformatics and Biostatistics Shared Resource.

