## Supplementary material for "*CDH1* loss remodels gene expression and lineage identity in human mammary epithelial cells": Supp. Fig.

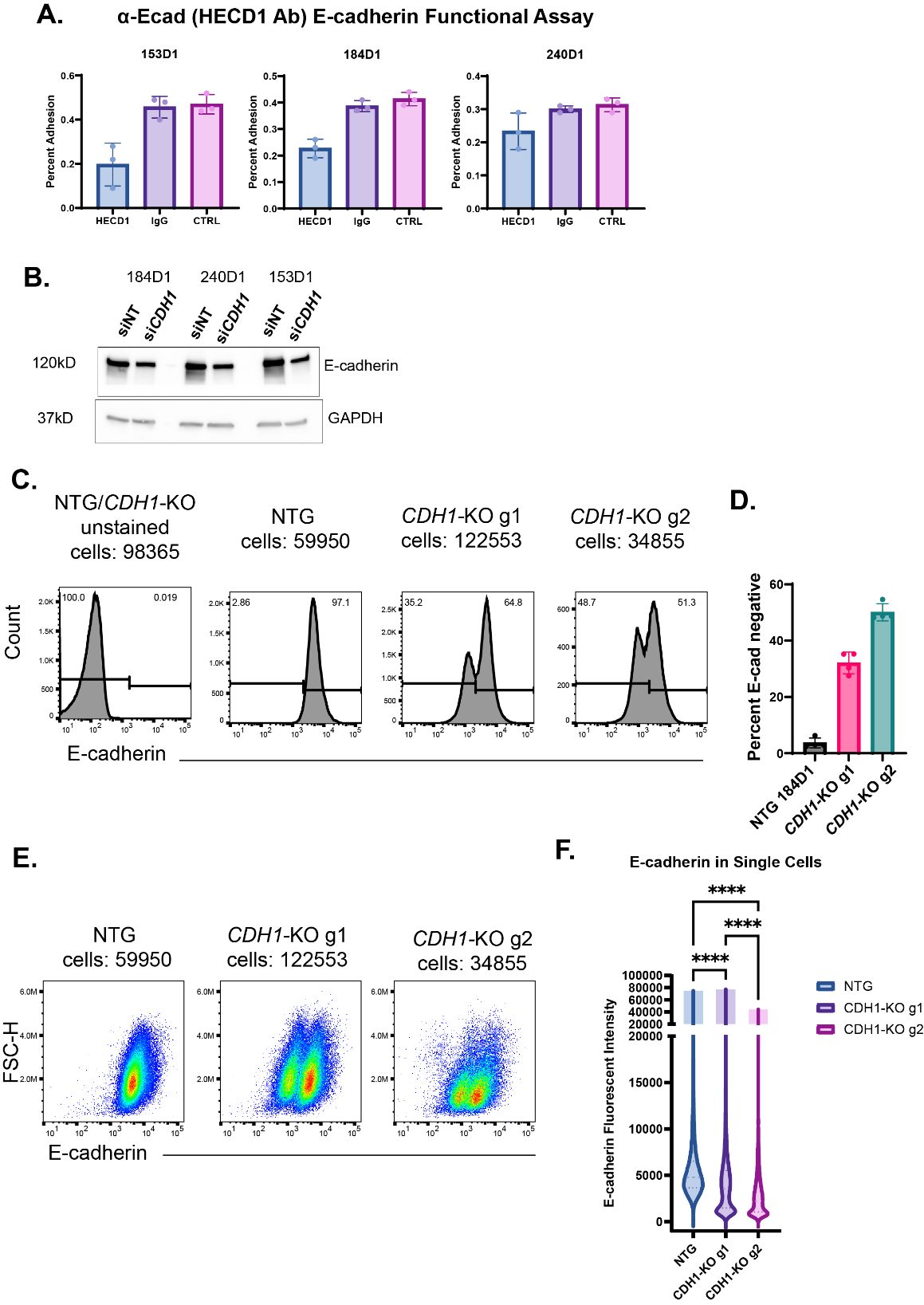

**Supplemental Figure 1. E-cadherin suppression within each model.** A) E-cadherin functional assay compares binding between HMEC 184D1, 153D1, 240D1 cells treated with HECD-1 vs. IgG and untreated. Plates were coated with E-cadherin at 3 ug/mL. Adhesion was quantified using GFP fluorescence. Bars represent average and standard deviation of percent adhesion (n=3). B) Western blot of E-cadherin abundance (clone: 24E10) in HMEC 184D1, 153D1, 240D1 cells treated with si*CDH1* or siNT for 72 hours. GAPDH was used as a loading control. C) Representative flow plots show *CDH1* KO efficiency. The first plot is HMEC 184D1 unstained negative control and was used to set the gate for E-cadherin negative (left of the gate) and E-cadherin positive (right of the gate). The next three plots were stained with E-cadherin Ab (clone: 24E10) conjugated to Alexa Fluor 647 (AF647). The x-axis is the logarithmic fluorescent intensity of AF647-E-cadherin. The percent of E-cadherin negative cells is in the top left of each plot and the percent of E-cadherin positive cells is in the top right of each plot. D) Quantification of percent E-cadherin negative cells across three biological replicates in both CDH1 KO g1 and g2 models and the control NTG model. E) Representative flow plots of size (FSC-H) vs. E-cadherin fluorescent intensity of individual cells in NTG, *CDH1*-KO g1 and *CDH1*-KO g2. F) Quantification of E. Violin plot showing the distribution of E-cadherin fluorescent intensity per cell in NTG, *CDH1*-KO g1 and *CDH1*-KO g2. **** adj. p < 0.0001 (one way-ANOVA).

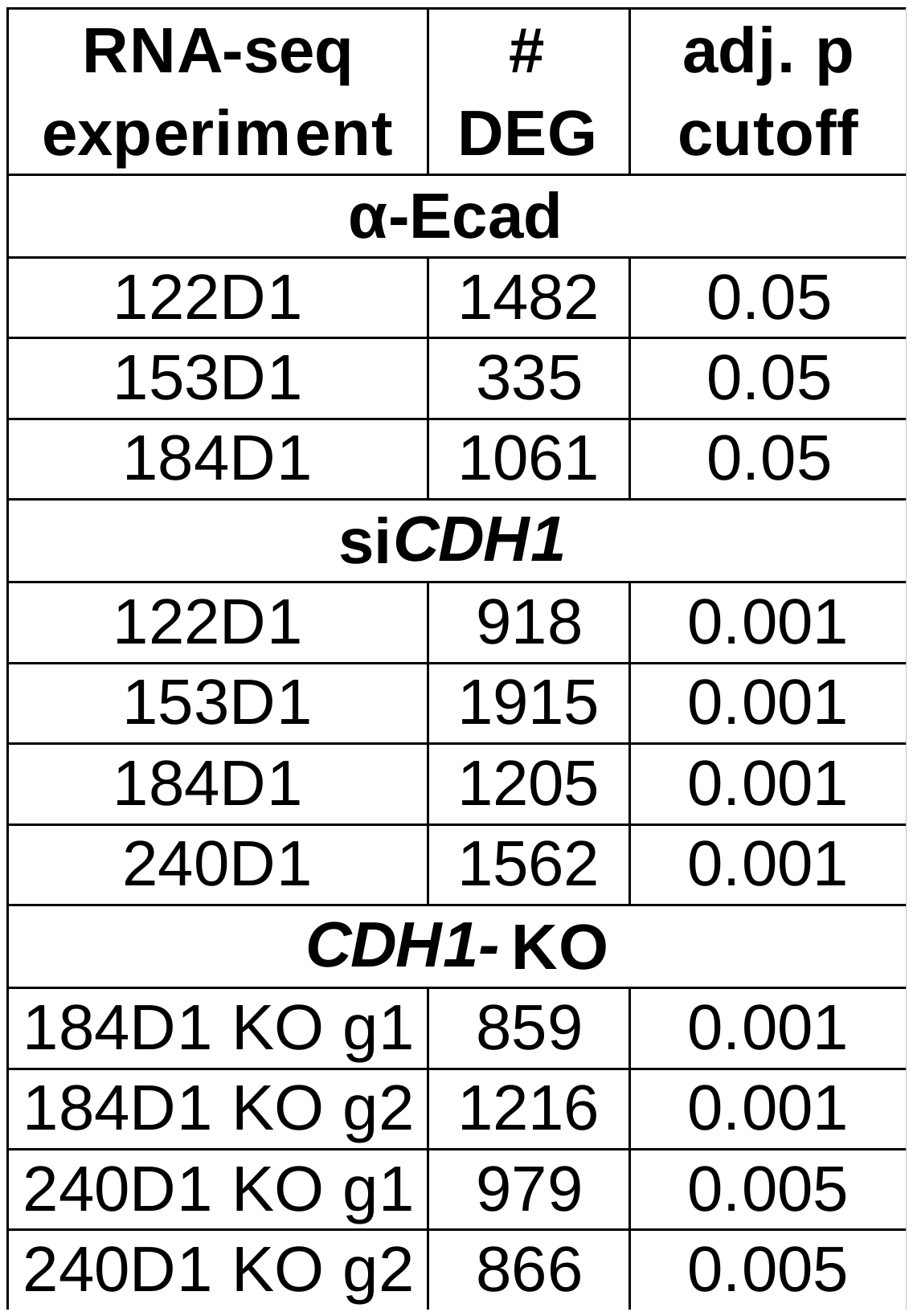

**Supplemental Table 1. Differentially expressed genes across all RNAseq datasets.** RNAseq datasets are organized by mode of E-cadherin suppression (α-Ecad, si*CDH1*, and *CDH1* KO) and cell model (122D1, 153D1, 184D1, 240D1). For each dataset the number of differentially expressed genes (DEG), and the adj. p value cutoff used to determine gene lists used for Fig. 1B-E are shown.

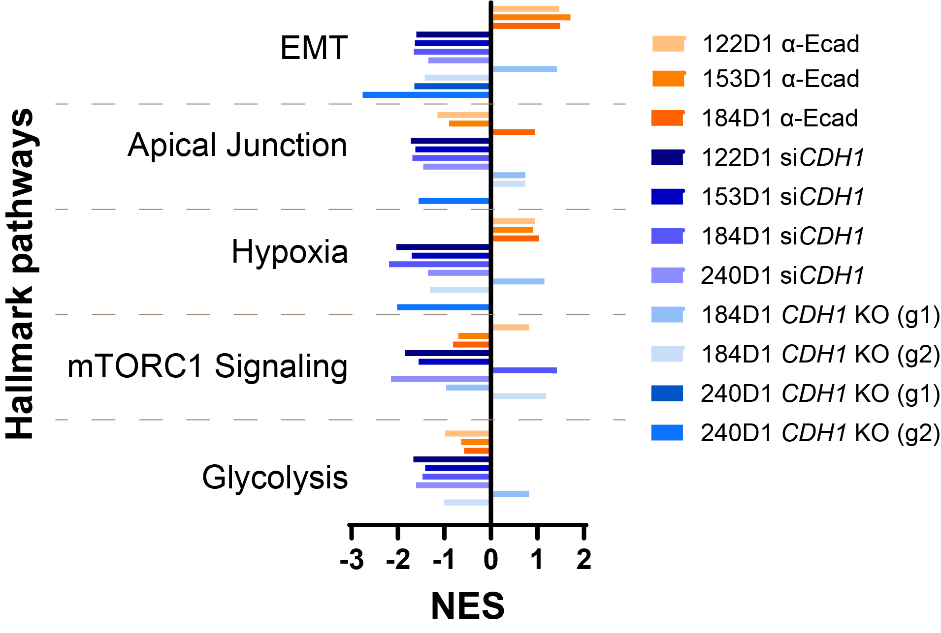

**Supplemental Figure 2. Fast gene set enrichment analysis** (FGSEA) normalized enrichment scores (NES) of Hallmark pathways (q value < 0.05; *EMT in 240D1 q val=0.06).

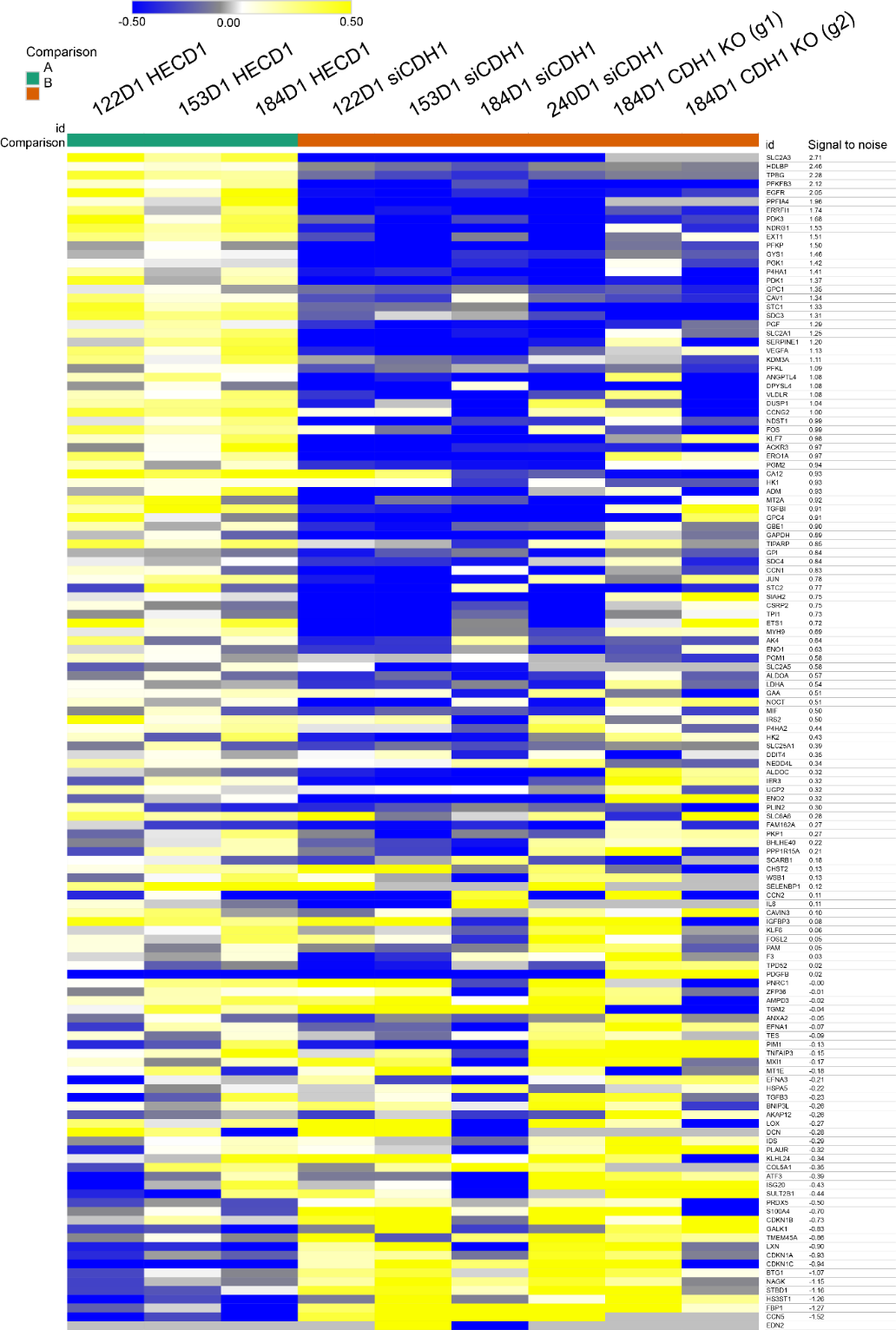

**Supplemental Figure 3. Differential gene expression within the Hypoxia signature.** Fold change of gene expression (p<0.05) between E-cadherin inhibition and genetic *CDH1* loss within each cell line (HMEC 122D1, 153D1, 184D1, 240D1) in the Hallmark pathway hypoxia signature.

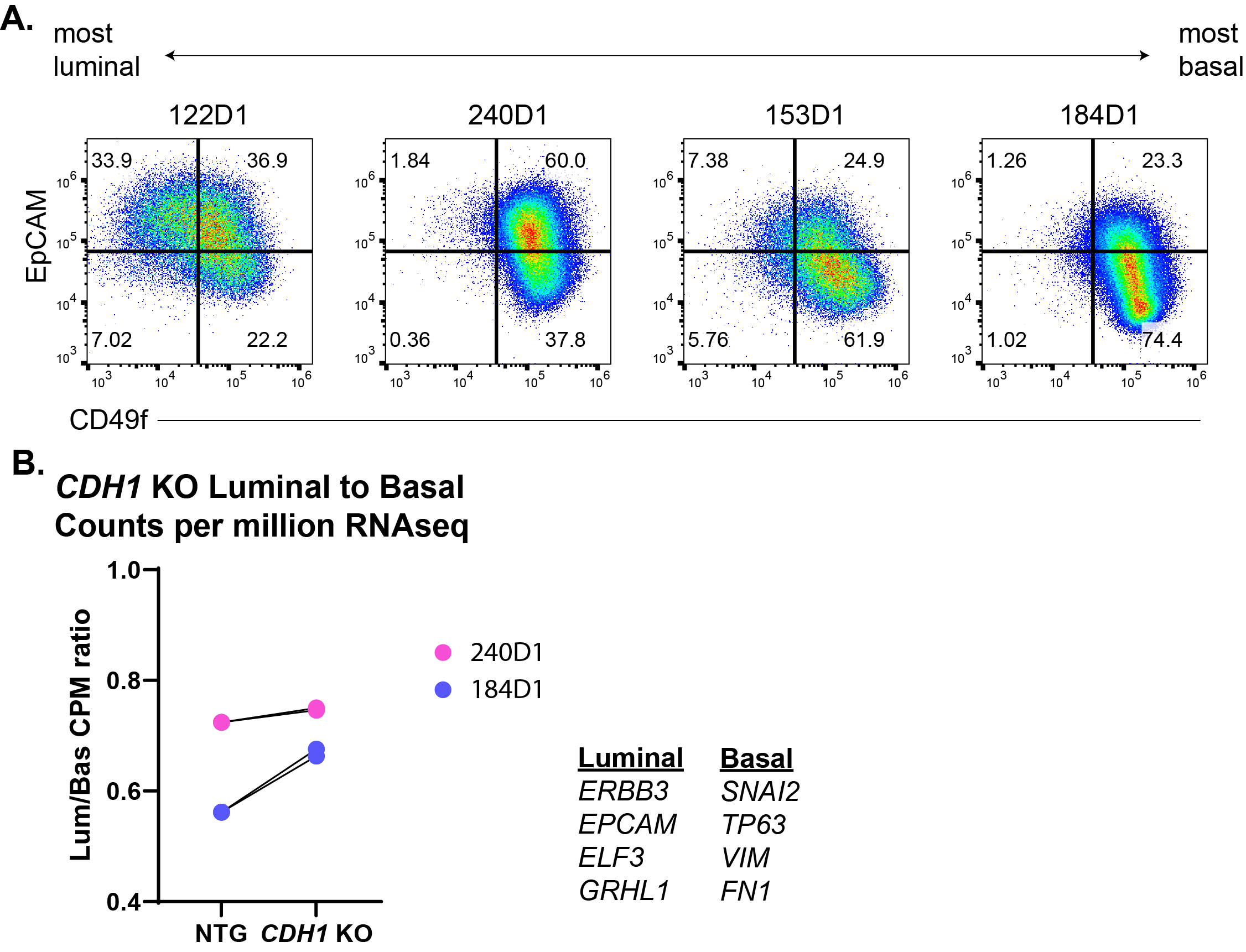

**Supplemental Figure 4. Luminal/Basal composition of HMEC models.** A) Distribution of luminal and basal cells within each of the 4 HMEC models using EpCAM and CD49f. B) Relative quantification of luminal/basal composition within HMEC *CDH1* KO models using counts per million (CPM) ratios of the luminal to basal genes highlighted in Figure 1D.

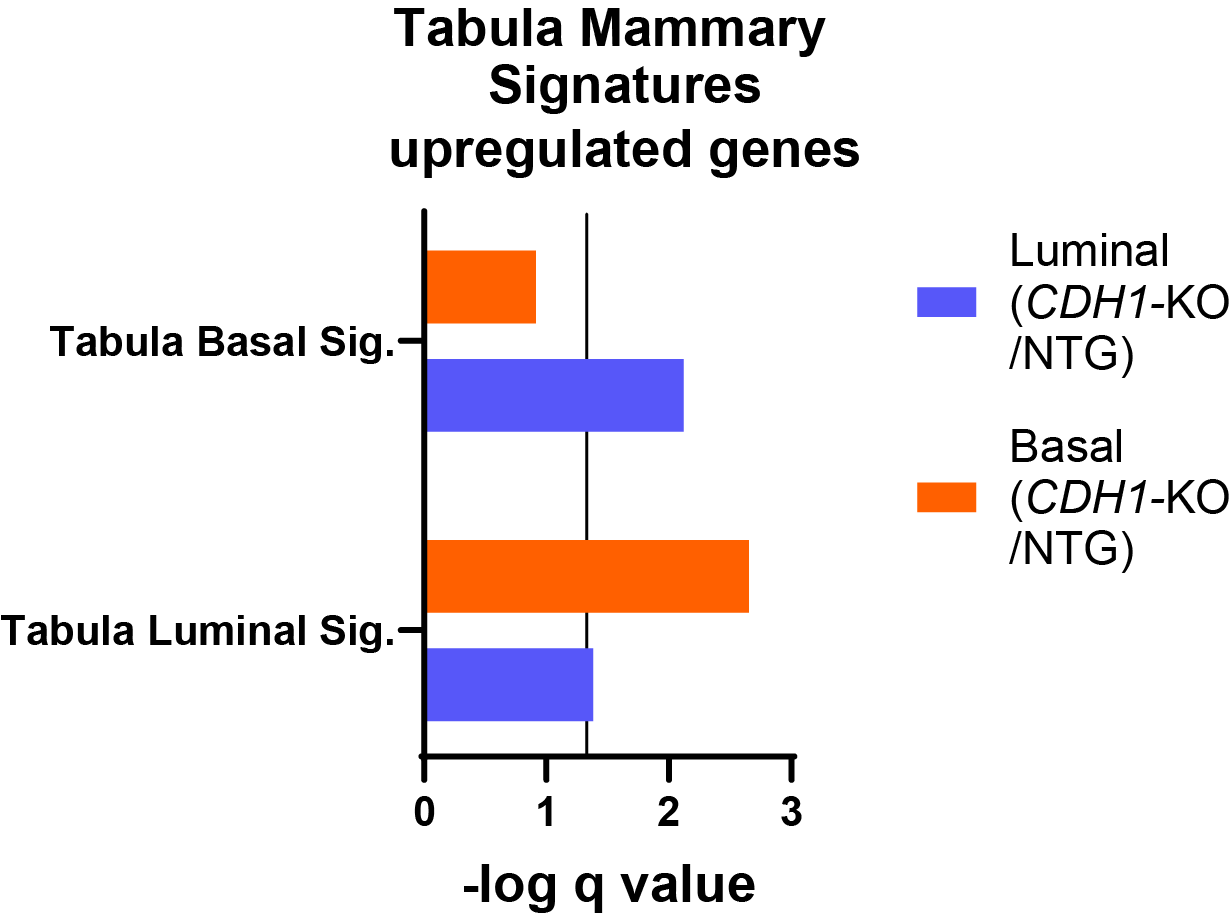

**Supplemental Figure 5. Tabula Muris gene signatures comparing luminal vs. basal in sc-RNAseq.** Overrepresentation analysis of only upregulated genes (luminal *CDH1*-KO vs. NTG; basal *CDH1*-KO vs. NTG) of mammary signatures in Tabula Muris cell-type pathways. (adj. p <1x10^-10^, -0.3>FC>0.3).

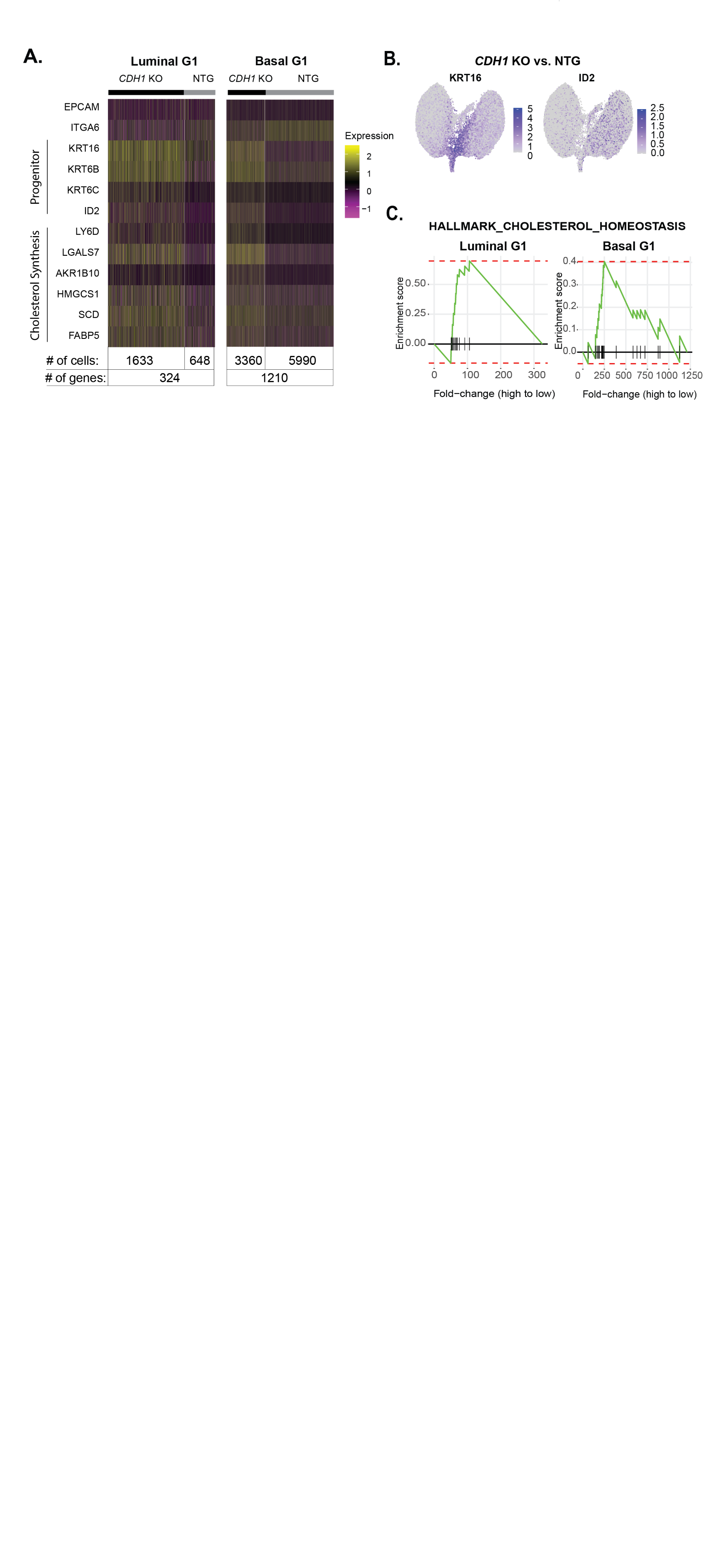

**Supplemental Figure 6. *CDH1* KO induces reprogramming in luminal and basal cells toward a luminal progenitor-like state.** A) Differentially expressed genes of *CDH1* KO vs. NTG cells within the luminal G1 and basal G1 populations relating to progenitor phenotype and cholesterol synthesis; tables on the right show the number of cells included in the analysis and the number of DGE for *CDH1* KO vs. NTG within Luminal G1 and Basal G1 cells, respectively (adj. p <1x10^-10^, -0.3>FC>0.3). B) FGSEA of Cholesterol Homeostasis Hallmark pathway in Luminal G1 and Basal G1 cells (luminal G1 NES= 2.48, adj. p=0.0027; basal G1 NES= 1.71, adj. p= 0.0706). C) Feature plots of two of the top 10 induced genes in *CDH1* KO luminal G1 and basal G1cells: KRT16 and ID2 mapped onto original sc-RNAseq feature plot including all NTG vs. *CDH1* KO cells (KRT16: avg. logFC= 1.79; pct.1=0.70; adj. p val = 0; ID2: avg. logFC= 2.03; pct.1=0.38; adj. p val = 0).

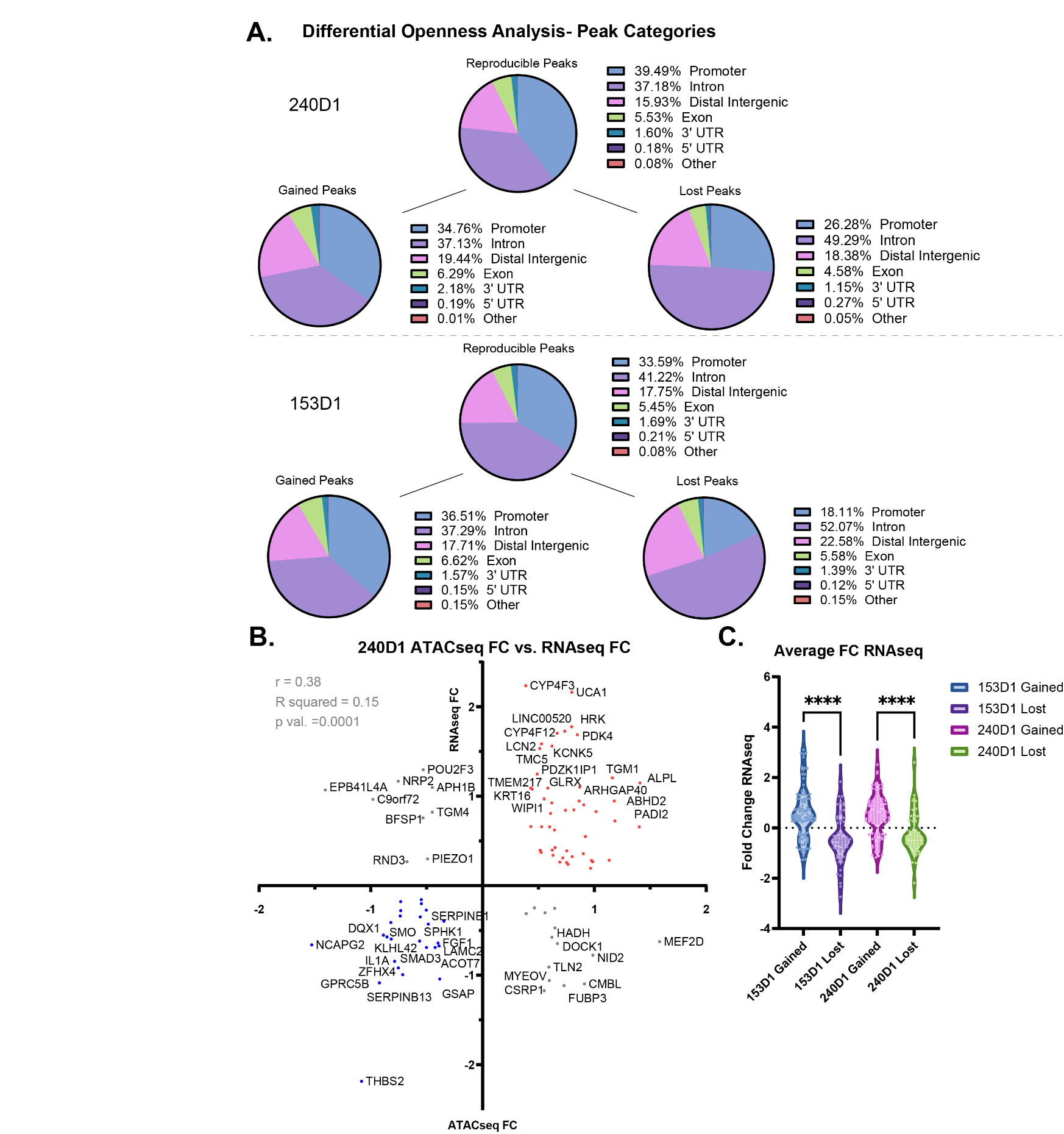

**Supplemental Figure 7. *CDH1*-KO chromatin landscape and associated gene expression changes.** A) Pie charts show the distribution of reproducible and differentially expressed peaks found in promoters, introns, distal intergenic regions, exons, 3’UTR, 5’UTR, and other. B) Scatter plot of 240D1 ATACseq fold changes of differential peaks vs. fold changes in corresponding gene expression from RNAseq. C) Violin plot shows fold changes of the differentially expressed genes from RNAseq (adj. p < 0.05) that are associated with gained or lost peaks within each cell line. The average fold change for genes associated with gained peaks in 153D1 was 0.55, whereas genes associated with lost peaks had an average fold change of -0.44. A similar trend was seen in the 240D1 (FC of genes associated with gained peaks = 0.46; FC of genes associated with lost peaks = -0.22). One-way ANOVA: **** (adj. p val < 0.0001).

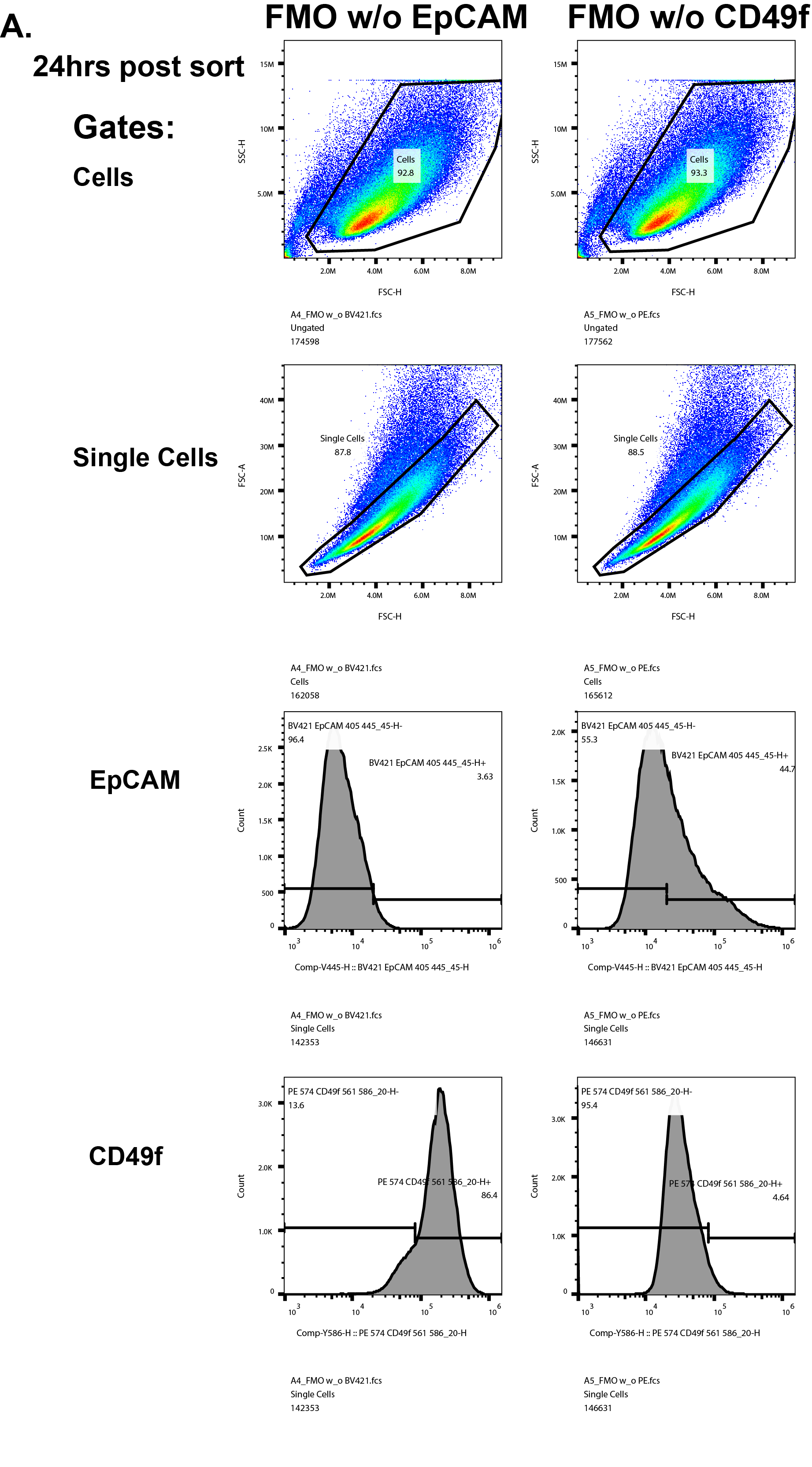

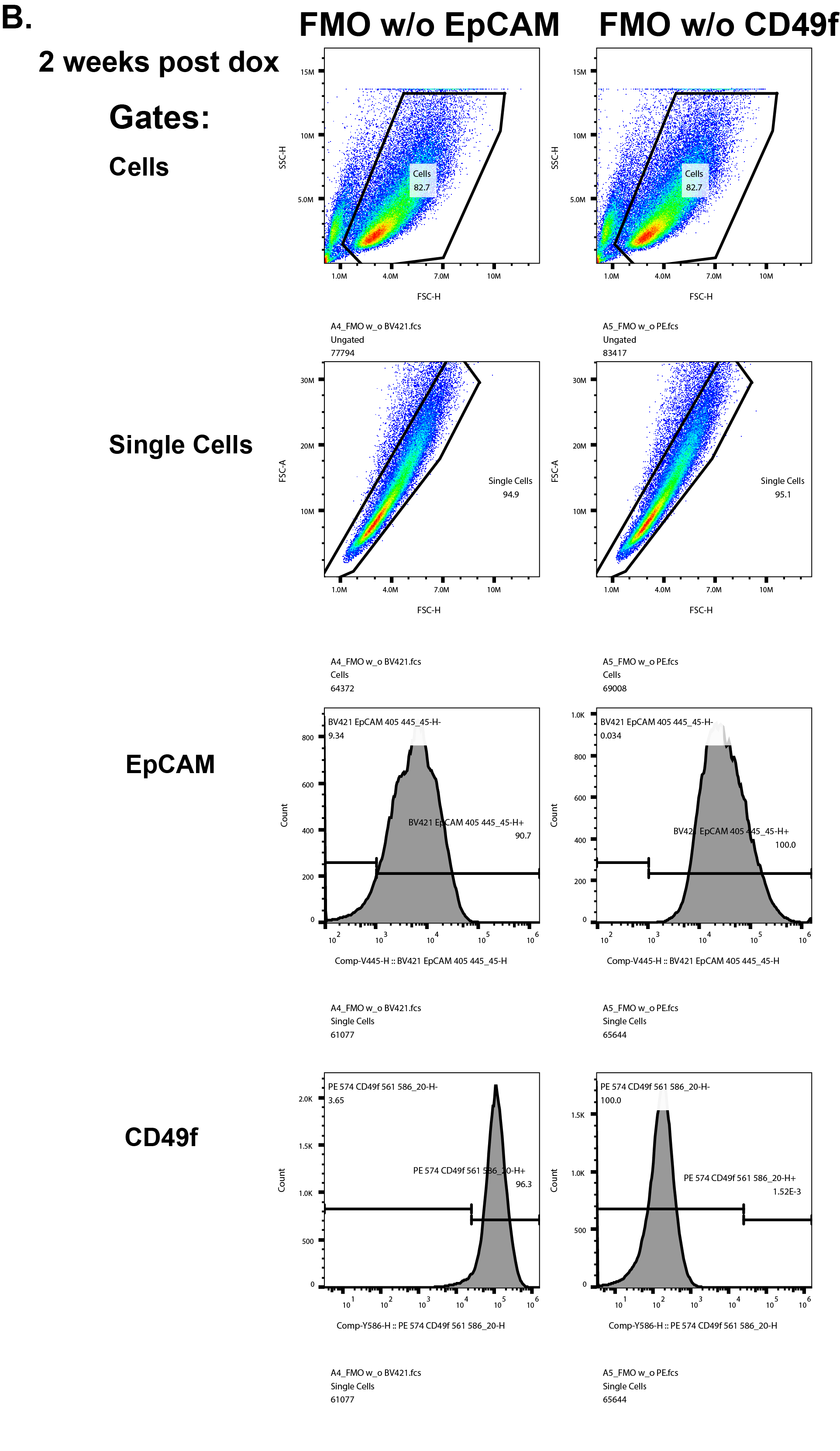

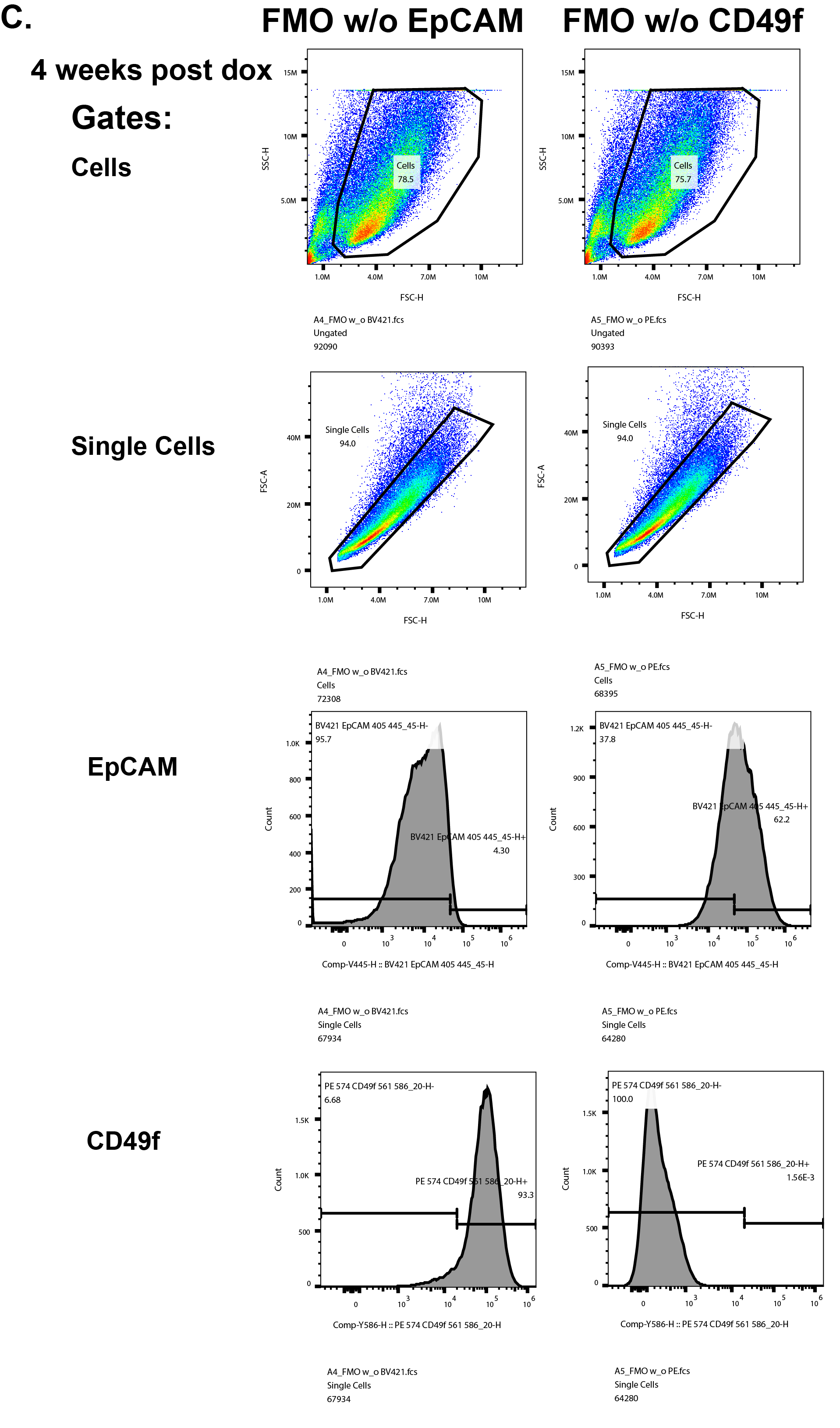

**Supplemental Figure 8. Gating for flow analyses after flow sort.** A-C) Fluorescence minus one (FMO) samples were used to set the gates for EpCAM and CD49f for all three post sort analyses including A) 24hrs post sort, B) 2 weeks post dox, and C) 4 weeks post dox. Gates for each day include Cells (SSC-H vs. FSC-H), Single Cells (FSC-A vs. FSC-H), EpCAM (Count vs. EpCAM-BV421), and CD49f (Count vs. CD49f-PE). The samples used to set the gates include a mixture of cells from each sorted population (NTG EpCAM Hi, NTG EpCAM Lo, *CDH1*-KO EpCAM Hi, and *CDH1*-KO EpCAM Lo). The left column shows the FMO without EpCAM antibody and the right column shows the FMO without the CD49f antibody. Antibody details can be found in methods.

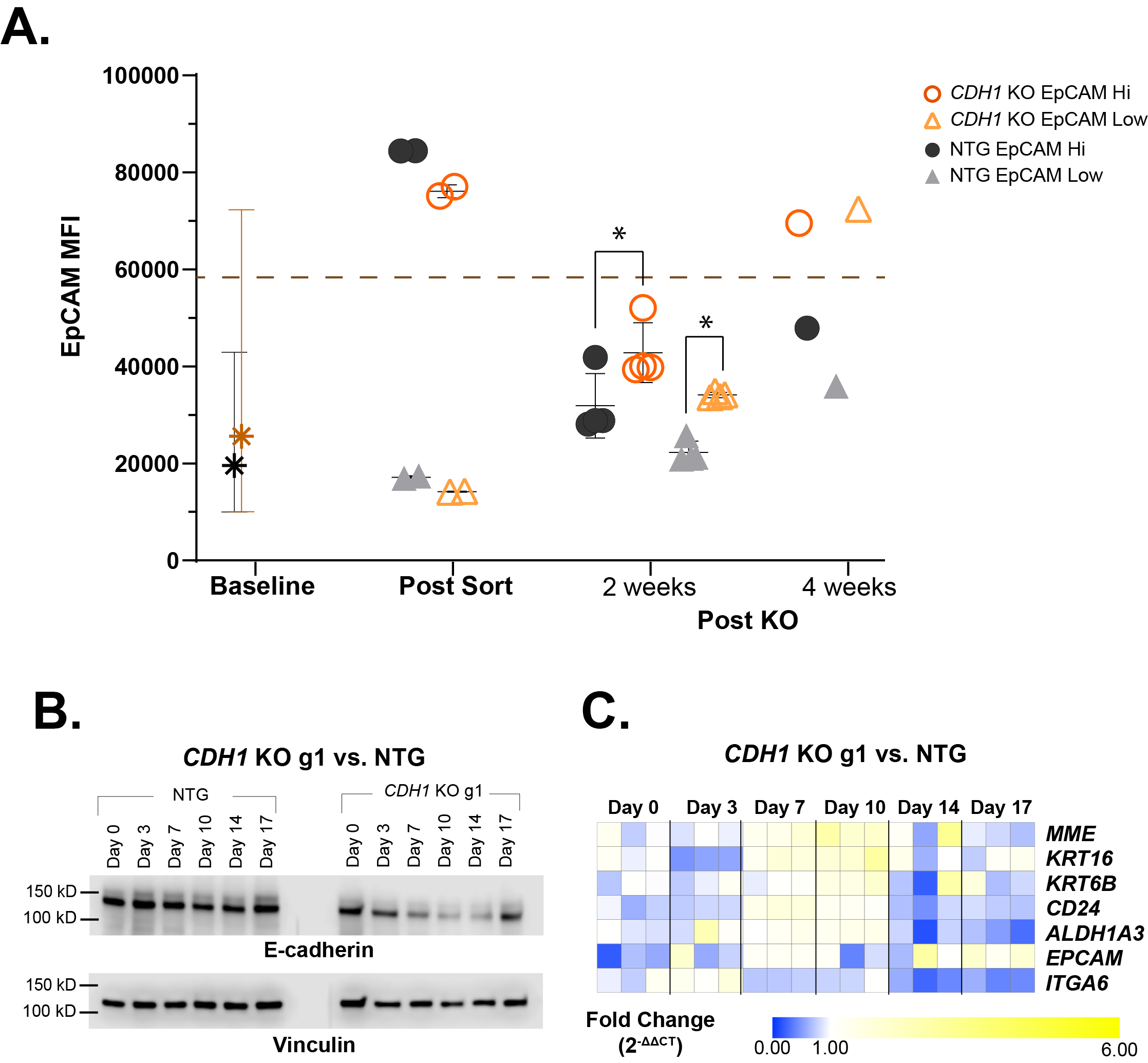

**Supplemental Figure 9. *CDH1*-KO Time Course with *CDH1*-KO g1.** A) EpCAM MFI (geomean) within parental/pre-sorted population (baseline) and each sorted population before induction of CDH1-KO (post-sort) and after *CDH1*-KO (post KO) both 2 week and 4 week time points. Baseline population error bars show 10% and 90% distribution of EpCAM MFI. (2 weeks, n =4, * adj. p val <0.05) B) Time course western blots of E-cadherin with Vinculin loading control. C) Paired time course samples analyzed via qPCR for basal and luminal gene expression.
