## Supplemental Methods for "*CDH1* loss remodels gene expression and lineage identity in human mammary epithelial cells"

***Single cell RNA-seq***

*Data Filtering and Processing*

The R package Seurat was utilized to process and analyze the scRNA sequencing data[1]. High quality cells were isolated via the criteria: number of transcripts mapped between 3000 and 45000, number of genes detected between 1500 and 8000, percent mitochondrial genes < 10%, and log10 genes per UMI > 0.8. Following filtering, 19,650 *CDH1*-KO g2 and 17,306 NTG cells remained included for downstream analysis. Feature counts were log normalized to correct for variations in sequencing depth between cells using the “NormalizeData” Seurat function and gene expression was transformed to z-scores using “ScaleData”.

The “FindVariableFeatures” function, using the variance-stabilizing transformation method, was applied to identify the top 3000 variable features. Principal component analysis was run on the variable features, using “RunPCA”, then the “RunUMAP” function was employed to perform the uniform manifold approximation and projection dimensional technique with 15 dimensions. Nearest neighbors were found using the “FindNeighbors” function with 15 dimensions, and clusters were determined using “FindClusters” at a resolution of 0.6.

*Cell Annotation*

Predicted cell cycle phases were determined using the “CellCycleScoring” function with the Seurat-defined “cc.genes” feature lists. Cells were classified as luminal or basal based on previously defined markers[2]. Using the “AddModuleScore” function, the transcriptome of each cell was compared with the luminal/basal marker lists and given a score for each phenotype. The lineage corresponding with the highest score was assigned to each cell.

*Clustering of Luminal Population (Supp. Fig. 6)*

The luminal cells were isolated into a separate Seurat object using the “subset” function and data was recentered to account for variability within only the luminal cells by running the “NormalizeData” and “ScaleData” functions. In parallel with the whole-data analysis, the top 3000 variable features were identified, PCA was done using the variable features, and UMAP was run with 15 dimensions. Cell neighbors were found, also using 15 dimensions, and clusters were defined with a resolution of 0.4.

*Clustering of Basal Population (Supp. Fig. 6)*

The basal cells were isolated into a separate Seurat object using the “subset” function and data was recentered to account for variability within only the luminal cells by running the “NormalizeData” and “ScaleData” functions. In parallel with the whole-data analysis, the top 3000 variable features were identified, PCA was done using the variable features, and UMAP was run with 15 dimensions. Cell neighbors were found, also using 15 dimensions, and clusters were defined with a resolution of 0.4.

*Differential Gene Expression Analysis*

Differentially expressed genes within each cluster were determined using the Seurat function “FindAllMarkers”, and comparisons between cell groups were calculated using the Wilcoxon rank sum test with the function “FindMarkers”. Fast gene set enrichment analysis (FSGEA) was used to identify Hallmark pathways of genes that passed these filters: -0.3>logFC >0.3, adj. p <1x10^-10^.

***ATAC-seq tools***

*Generate Heatmap- Galaxy* (<https://usegalaxy.org/>)

1. Compute matrix
   a. Select regions (3 regions: maintained broad peaks, lost broad peaks, gained broad peaks BED files)
   b. Score files (4: upload bigwig files for all samples; siNT_A, siNT_B, siCDH1_A, siCDH1_B; per cell line)
   c. Specify new labels (siNT_A siNT_B siCDH1_A siCDH1_B)
   d. computeMatrix has two main output options
      i. Choose reference-point
      ii. Choose center of region
   e. Distance upstream and downstream are set to default of 1000bp
   f. Sort regions
      i. Choose descending order
   g. All other options are set to the default options
2. plotHeatmap
   a. Matrix file is the "computeMatrix" file created in step 1
   b. Advanced options
      i. Sort regions: choose descending order
      ii. Missing data color: choose black
      iii. List of colors for each heatmap.: Enter- white,blue white,blue
      iv. Title of plot: Cell line
      v. Did you use multiple groups of regions: choose Yes
      vi. Any other options were left as the default setting

*Peak Overlap - Galaxy* (<https://usegalaxy.org/>)

- Used gained and lost peak files from reproducible peaks analysis
  - 240D1 diff open analysis_gained peaks
  - 240D1 diff open analysis_lost peaks
  - 153D1 diff open analysis_gained peaks
  - 153D1 diff open analysis_lost peaks

1. bedtools Intersect intervals:
   a. Shared gained/lost peaks between 240D1 and 153D1
      i. File A	240D1 diff open analysis_gained/lost peaks
      ii. File B	153D1 diff open analysis_gained/lost peaks
   b. Combined or separate output files: single output containing intersections of any 'input B' lines with A
   c. Calculation based on strandedness?: overlaps on either strand
   d. What should be written in the output file?: Write the original A and B entries plus the number of base pairs of overlap between the two features. Only A features with overlap are reported. Restricted by the fraction and reciprocal option (-wo)
   e. Required overlap: 0.4	-40%
   f. Require that the fraction of overlap be reciprocal for A and B: Yes
   g. The rest of the parameters were set to the default

*Peak gene associations- Cistrome* **(http://cistrome.org/)**

1. BETA minus
   a. Use shared gained/lost peaks file created above
   b. Distance from TSS: 30kb
   c. Take resulting gene peak associations and make export to excel
   d. Remove duplicate gene names
   e. This will give you the total number of genes for the shared peaks
2. BETA basic
   a. Take shared peak files from above (not output of BETA minus)
   b. Take RNAseq DGE (adj. p val < 0.05)
   c. Compare upregulated genes associated with gained peaks from 153D1 and 240D1
   d. Compare downregulated genes associated with lost peaks

***Immunoblotting***

Whole-cell lysates were obtained by incubating cells in RPPA lysis buffer for 45′ on ice. Cells were centrifuged at ~16,000× *g* for 15 m at 4 °C and the resulting supernatant was collected for analysis. Protein concentrations were measured and normalized using the Pierce BCA protein assay kit (#23225). Protein loading was kept consistent by mass across matched experiments, and standard methods were used to perform SDS-PAGE. Proteins were transferred onto PVDF membranes. Antibodies were used according to manufacturer’s recommendations: *CDH1* (Cell Signaling, CAT# 3195S, clone #24E10) and GAPDH (Cell Signaling, CAT #5174, clone #D16H11 . Secondary antibodies were used according to manufacturer’s instruction and were obtained from Jackson ImmunoResearch Laboratories (West Grove, PA, USA), goat anti-mouse IgG (cat# 115-035-068) and goat anti-rabbit IgG (cat# 111-035-045). Chemiluminescence was used to detect antibodies and the LICOR c-Digit (LI-COR Biosciences, Lincoln, NE, USA) was used to develop immunoblots. GAPDH served as a loading control.

***E-cadherin functional assay***

Untreated 96-well plates were coated with recombinant human E-cadherin (R&D Systems #8505-EC) (3ug/mL) for overnight incubation at 4C. Untreated wells (Dulbecco’s phosphate-buffered saline (DPBS) - coating buffer only) were used as negative controls. The next morning, wells were aspirated and plates were incubated with fresh blocking buffer (1% BSA in DPBS) at 37C for 1 hour. During this time, cells were dissociated, blocked, and treated with HECD-1 (25ug/mL), IgG (25ug/mL), or untreated. Cells were treated for 4 hours at 37C. Since our cells express GFP, we used GFP fluorescence as a measure of relative number of cells. A fluorescent plate reader was used to measure fluorescence after the 4 hour incubation (excitation: 488nm; emission: 507nm). Cells were washed with PBS and then read on the fluorescent plate reader again. Percent adhesion was calculated by the following equation. % adhesion = (OD after wash)/OD before wash)*100. Average OD was used among technical triplicates (n=3).

***ATACseq analysis additional references***

1. Andrews, S. (2010). *FastQC: a quality control tool for high throughput sequence data*. Available online at: [http://www.bioinformatics.babraham.ac.uk/projects/fastqc](https://nam02.safelinks.protection.outlook.com/?url=http%3A%2F%2Fwww.bioinformatics.babraham.ac.uk%2Fprojects%2Ffastqc&data=05%7C02%7CMARGARET.MUSICK%40CUANSCHUTZ.EDU%7Cea92a176176f4936e07f08deb961a096%7C563337caa517421aaae01aa5b414fd7f%7C0%7C0%7C639152026133531873%7CUnknown%7CTWFpbGZsb3d8eyJFbXB0eU1hcGkiOnRydWUsIlYiOiIwLjAuMDAwMCIsIlAiOiJXaW4zMiIsIkFOIjoiTWFpbCIsIldUIjoyfQ%3D%3D%7C0%7C%7C%7C&sdata=oUBtJPLCs0iaakt2TwqdxLhbq2qw%2BZ7DXvzjPiRuAGQ%3D&reserved=0)
2. Ewels, P., Magnusson, M., Lundin, S., & Käller, M. (2016). MultiQC: summarize analysis results for multiple tools and samples in a single report. *Bioinformatics*, 32(19), 3047–3048. PMID: 27312411
3. Wingett, S. W., & Andrews, S. (2018). FastQ Screen: A tool for multi-genome mapping and quality control. *F1000Research*, 7, 1338. PMID: 30254741
4. Langmead, B., & Salzberg, S. L. (2012). Fast gapped-read alignment with Bowtie 2. *Nature Methods*, 9(4), 357–359. PMID: 22388286
5. Li, H., Handsaker, B., Wysoker, A., Fennell, T., Ruan, J., Homer, N., Marth, G., Abecasis, G., Durbin, R., & 1000 Genome Project Data Processing Subgroup. (2009). The Sequence Alignment/Map format and SAMtools. *Bioinformatics*, 25(16), 2078–2079. PMID: 19505943
6. Quinlan, A. R., & Hall, I. M. (2010). BEDTools: a flexible suite of utilities for comparing genomic features. *Bioinformatics*, 26(6), 841–842. PMID: 20110278
7. Ramírez, F., Ryan, D. P., Grüning, B., Bhardwaj, V., Kilpert, F., Richter, A. S., Heyne, S., Dündar, F., & Manke, T. (2016). deepTools2: a next generation web server for deep-sequencing data analysis. *Nucleic Acids Research*, 44(W1), W160–W165. PMID: 27079975
8. Zhang, Y., Liu, T., Meyer, C. A., Eeckhoute, J., Johnson, D. S., Bernstein, B. E., Nusbaum, C., Myers, R. M., Brown, M., Li, W., & Liu, X. S. (2008). Model-based analysis of ChIP-Seq (MACS). *Genome Biology*, 9(9), R137. PMID: 18798982
9. For IGV:
   Robinson, J. T., Thorvaldsdóttir, H., Winckler, W., Guttman, M., Lander, E. S., Getz, G., & Mesirov, J. P. (2011). Integrative Genomics Viewer. *Nature Biotechnology*, 29(1), 24–26. PMID: 21221095
10. For DiffBind:
    Ross-Innes, C. S., Stark, R., Teschendorff, A. E., Holmes, K. A., Ali, H. R., Dunning, M. J., Brown, G. D., Gojis, O., Ellis, I. O., Green, A. R., Ali, S., Chin, S. F., Palmieri, C., Caldas, C., & Carroll, J. S. (2012). Differential oestrogen receptor binding is associated with clinical outcome in breast cancer. *Nature*, 481(7381), 389–393. PMID: 22217937

**KEY RESOURCES TABLE**

| **REAGENT OR RESOURCE** | **SOURCE** | **IDENTIFIER** |
| --- | --- | --- |
| **Antibodies** | | |
| Estrogen Receptor alpha (6F11) | Leica | Cat # ER-6F11-L-F |
| HECD-1 IgG mouse monoclonal (2mg/mL) | Thermo Fisher | Cat # 13-1700;  RRID: AB_2533003 |
| Chrome Pure Mouse, whole IgG | Jackson ImmunoResearch | Cat # 015-000-003;  RRID: AB_2337188 |
| *CDH1* IgG rabbit (clone # 24E10) | Cell Signaling | Cat # 3195S |
| Goat anti-rabbit IgG | Jackson ImmunoResearch | Cat # 111-035-045;  RRID: AB_2337938 |
| Goat anti-mouse IgG | Jackson ImmunoResearch | Cat # 115-035-068;  RRID: AB_2338505 |
| EpCAM-BV421 (clone 9C4, mouse IgG2b, 1:200) | Biolegend | Cat #324220  RRID:AB_2563847 |
| CD10-APC/Fire 750 (clone HI10a, mouse IgG1, 1:200) | Biolegend | Cat #312230  RRID:AB_2616719 |
| CD49f-PE (clone GoH3, Rat IgG2a, 1:400) | Biolegend | Cat #313612  RRID:AB_893373 |
| E-cadherin- AF647 (clone 24E10, Rabbit IgG, 1:200) | Cell Signaling | Cat #9835  RRID:AB_10828228 |
| **Experimental models: Cell lines** | | |
| HMEC 184D-D1L | Dr. Martha Stampfer, LBNL |  |
| HMEC 153L-D1 | Dr. Martha Stampfer, LBNL |  |
| HMEC 122L-D1 | Dr. Martha Stampfer, LBNL |  |
| HMEC 240LB-D1 | Dr. Martha Stampfer, LBNL |  |
| **Chemicals, Peptides, and Recombinant Proteins** | | |
| MEBM | Lonza | Cat #cc-3151 |
| 5.0ug/mL Insulin | Thermo Fisher | Cat # 41400-045 |
| 70.0ug/mL bovine pituitary extract | Thermo Fisher | Cat # 13028014 |
| hydrocortisone | Sigma-Aldrich | Ca t# H4001-1G |
| EGF | Thermo Fisher | Cat # 236EG200 |
| transferrin | Invitrogen | Cat # 11108-016 |
| Isoproterenol (IP) | Sigma-Aldrich | Cat # I5627 |
| glutamine | Sigma-Aldrich | Cat # 59202C |
| DMEM/F12 | Corning | Cat # 10-092-CV |
| BSA | Sigma-Aldrich | Cat # A4161 |
| Cholera Toxin | Sigma-Aldrich | Cat # C8052 |
| Oxytocin | Sigma-Aldrich | Cat # O6379-1MG |
| Albumax | Invitrogen | Cat # 11020-021 |
| Estradiol (E2) was from Sigma | Sigma-Aldrich | Cat # E2758 |
| Hoechst 33258 | Thermo Fisher | Cat # 62249 |
| DNA Extraction Buffer | Cell Signaling | Cat # 42015 |
| Proteinase K | Cell Signaling | Cat # 10012 |
| RNase A | Cell Signaling | Cat # 7013 |
| **Critical Commercial Assays** | | |
| Illumina Single Cell 3' RNA Prep | Illumina |  |
| ATAC-seq Kit | Active Motif | Cat # 53150 |
| ATAC-Seq Spike-In Control | Active Motif | Cat # 53154 |
| DNA Library Prep Kit for Illumina® | Active Motif | Cat # 53220 |
| Dual Index Primers Set 1 for Illumina® | Active Motif | Cat # 53221 |
| PowerUp SYBR Green Master Mix | Life Technologies, Inc. | Cat # 100029284 |
| iSCRIPT cDNA synthesis kit | BioRad | Cat # 1708890 |
| RNeasy mini kit | Qiagen | Cat # 74034 |
| LIVE/DEAD™ Fixable Blue Dead Cell Stain Kit, for UV excitation | Thermo Fisher | Cat # L23105 |
| **Oligonucleotides** | | |
| RPL30 (exon 3) forward primer: 5’ GTC CTG GGG TAC AAG CAG AC 3’ | Sikora Laboratory | N/A |
| RPL30 (exon 3) reverse primer: 5’ CTG GGC AGT TGT TAG CGA GA 3’ | Sikora Laboratory | N/A |
| **Software and Algorithms** | | |
| Enrichr | Maayan Lab | <https://maayanlab.cloud/Enrichr/>  RRID:SCR_001575 |
| Morpheus | Broad Institute | <https://software.broadinstitute.org/morpheus/> |
| GraphPad Prism 9-10 | GraphPad | <https://www.graphpad.com/>  RRID: SCR_002798 |
| BBDuk | R Studio | RRID:SCR_016969 |
| STAR (2.6.0a) | R Studio | RRID:SCR_004463 |
| edgeR package | R Studio | RRID:SCR_012802 |
| limma R package | R Studio | RRID:SCR_010943 |
| Profiler R | R Studio | RRID:SCR_016884 |
| Molecular Signatures Database | R Studio | RRID:SCR_016863 |
| CU Anschutz Bioinformatics RNAseq Analysis tool | CU Anschutz Bioinformatics Core | [https://bioinformatics.cuanschutz.edu/UC_RNA-seq](https://bioinformatics.cuanschutz.edu/rnaseq) |
| Venny 2.1 (Venn Diagrams) | CNB | <https://bioinfogp.cnb.csic.es/tools/venny/> |
| **Others** | | |
| Countess 2 Automated Cell Counter | Invitrogen | Cat # AMQAX1000 |
| 4200 TapeStation System | Agilent | Cat # G2991BA |
| Qubit 4 Fluorometer | ThermoFisher | Cat # Q33238 |
| NovaSeq 6000 | Illumina | Cat # 20012850 |
| Magnetic Separation Rack, 0.2 mL Tubes | EpiCypher | Cat # 10-0008 |
